# Proteotype Co-evolution and Diversity in Mammals

**DOI:** 10.1101/2022.06.08.495293

**Authors:** Qian Ba, Yuanyuan Hei, Anasuya Dighe, Wenxue Li, Jamie Maziarz, Irene Pak, Shisheng Wang, Günter P. Wagner, Yansheng Liu

## Abstract

Evolutionary profiling has been largely limited to the nucleotide level. Using consistent proteomic methods, we quantified proteomic and phosphoproteomic layers in fibroblasts from 11 common mammalian species, with transcriptomic variability as reference. The co-variation analysis indicates that transcript and protein expression robustness across mammals remarkably follows functional role, with extracellular matrix-associated expressions being most variable, demonstrating strong transcriptome-proteome co-evolution. Interestingly, the variability control of gene expression is universal at both inter-individual and inter-species scales, but of different extent. RNA metabolism processes particularly show the higher inter-species versus inter-individual variations. Our results further uncover that while ubiquitin-proteasome system is extremely conserved in mammals, the lysosome-mediated protein degradation exhibits a remarkable variation between mammalian lineages. Additionally, the phosphosite profiles reveals phosphorylation co-evolution network independent of protein abundance.

## Introduction

Despite the scalable nucleotide sequencing performed in evolutionary biology, it is ultimately protein abundances and activities that, to a large part, define the organism phenotype. Recently, a qualitative proteome landscape for 100 taxonomically diverse organisms was established by mass spectrometry (MS)-based analysis (*1*). However, a quantitative evolutionary comparison of emerged proteomes across multiple species, such as mammals represents, so far, uncharted territory. Ribosome-profiling (Ribo-seq) was used as a proxy for quantifying proteins synthesized, which discovered co-evolutionary patterns across transcriptome and translatome in five mammals (*2*). However, both the proteome dynamic range and protein degradation cannot be directly measured by Ribo-seq. On the other hand, the current reproducible proteomics workflows exemplified by data-independent acquisition (DIA) MS has achieved a favorable reproducibility and quantitative performance for the global proteome (*3, 4*), with data quality thoroughly and widely assessed (*5-8*). Yet, a systematic, unbiased multi-species quantitative effort has been lacking to link individual variability to species level variability (*9*) and to understand phosphorylation signaling among multiple species in a comprehensive manner.

## Results

### Steady-state and diverse proteotype across 11 mammalian species

Proteotype is defined as the proteome complement of a genotype (*10, 11*). To understand the functional and molecular basis of proteotype evolution in mammals, we profiled the steady-state proteomes and phosphoproteomes of skin fibroblast cells from 11 common mammalian species (**Figure 1a**) by DIA-MS (*12, 13*). Considering that different cell types present a major variable factor in profiling gene expression (*14-16*), we herein exclusively analyzed the fibroblast cells commonly used for evolutionary studies. The mammalian species we analyzed represent two major phylogenetic clades, *Euarchontoglires* (EAOG: primates, rodents and their relatives) and *Laurasiatheria* (LAUT: carnivors, hoofed animals and their relatives), together with an evolutionarily distant species, Opossum (*Monodelphis domestica*), as the outgroup. The EAOG clade included Rabbit (*Oryctolagus cuniculus*), Rat (*Rattus norvegicus*), Monkey (*Mucaca mulatta*), and Human (*Homo sapiens*), whereas the LAUT clade included Sheep (*Ovis aries*), Cow (*Bos taurus*), Pig (*Sus scrofa*), Dog (*Canis lupus*), Cat (*Felis catus*), and Horse (*Equus caballus*).

**Fig. 1.**
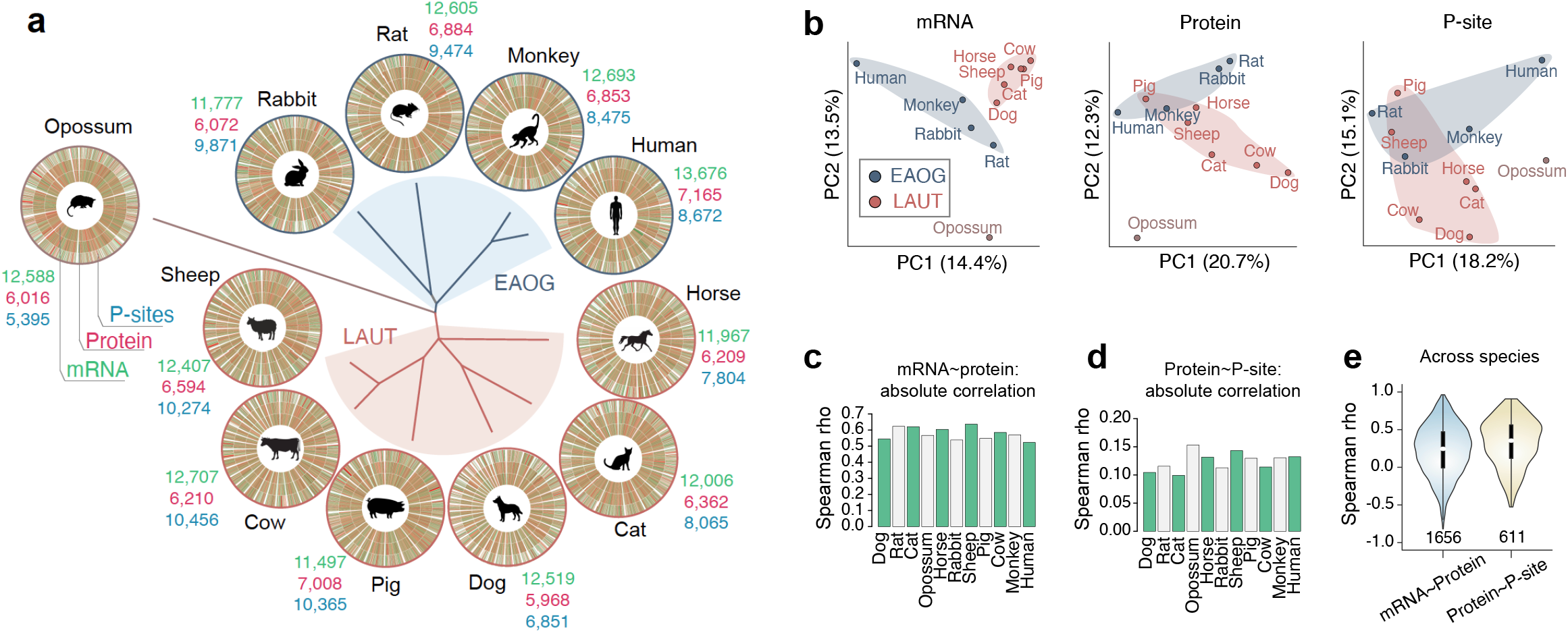
Identification and quantification overview of transcriptome, proteome and phosphoproteome across 11 mammalian species. **(a)** Phylogenetic relationships among 11 mammalian species including 10 Boreotherian mammals and the opossum *Monodelphis domestica* as an outgroup. In Boreotherian mammals, 6 Laurasiatheria (LAUT) were colored as red and 4 Euarchontoglires (EAOG) were colored as blue. The transcriptome, proteome, and phosphoproteome in skin fibroblast cells of all these species were shown in circos plots. The identification numbers of mRNA (TPM>1), protein or phosphorylation sites (P-sites) in individual species are shown in green, red and blue colors. **(b)** Principal component analysis of mRNA, protein and P-site profiles in 11 species. (**c-d**) Within-species correlation between different layers (mRNA∼protein (**c**) and Protein∼P-site (**d**)) of all genes/sites in individual species. Spearman’s *rho* for every species was individually calculated from all detected genes/sites in single species. (**e**) Gene-specific and cross-species correlation between different layers (mRNA∼protein and Protein∼P-site). Spearman’s *rho* for every gene/site was individually calculated from the data set of 11 species. All *rho* values were then summarized as violin plots.

Our spectral library-free DIA-MS (*4, 17*) was able to detect an average of 6,490 protein groups (peptide-and protein-FDR <1%) (*18*) in the fibroblasts of different species, ranging from 5,968 (dog) to 7,165 (human) identified proteins. In most species, our phosphoproteomic analysis measured >8,000 unique, confidently localized phosphosites (P-sites). In addition, we profiled the mRNA levels for an average of 12,400 genes in all the samples by mRNA sequencing (RNA-seq) to study possible post-transcriptional regulations (**Figure 1a**). After one-to-one pairwise gene ortholog mapping (*19*) (see **Methods**), 4,353 transcripts (Transcripts Per Million transcripts, TPM>1), 1,660 proteins, and 546 phosphoproteins (containing 611 P-sites) were filtered to be overlapping across species (**Figure S1a**). All the individual replicates clustered together in the hierarchical clustering analysis (HCA) based on both mRNA and protein quantities (Pearson R=0.9822 and 0.9855, **Figure S1b-c**). Together, our datasets enabled a deep and precise quantification of proteotype and co-varying signaling at each molecular layer (**Figure S2a**) in mammals.

We subsequently explored the extent of regulation at different molecular layers. According to principal component analysis (PCA) (**Figure 1b**), we found that, compared to the mRNA data, the proteome and phosphoproteome data both showed a smaller power in separating EAOG and LAUT (90 Myr ago), indicating substantial proteotype variability. The transcriptome and proteome of Opossum (160 Myr) are both quantitatively distant from EAOG and LAUT species, as expected. Furthermore, in the absolute scale, within each species the mRNAs fairly predicted protein levels (Spearman *rho is* 0.50-0.65, **Figure 1c**) (*20-22*), whereas the absolute protein abundances only poorly predict the P-site intensities (*rho*=0.10-0.15, **Figure 1d**). Notably, the relative, cross-species mRNA∼protein correlation is centered at Spearman’s *rho* of merely 0.224, based on all quantified mRNA∼protein pairs across species (n=1,656, **Figure 1e**), arguing pervasive protein level remodeling between species. Interestingly, due to the removal of detectability bias among P-sites, the cross-species protein∼P-site correlation is higher than the within-species comparison, but still weak for many P-sites (mean of *rho* for all P-sites, 0.326). Furthermore, the *rho* distributions for mRNA∼protein and Protein∼P-site tend to be gene function class-dependent (**Figure S2b-d**). Therefore, the post-transcriptional and phosphorylation-mediated regulations among mammals are both extensive and gene function-specific.

### Transcriptome-proteome co-evolution across gene classes

To associate mRNA and protein levels with evolutionary process, we firstly performed the phylogenetically independent contrast (PIC) analysis (*23*) (see **Methods** and **Figure S3**). The thus determined mRNA-and protein-PICs across phylogenetical nodes are positively correlated (averaged R= 0.306, **Figure S3a-b**), confirming strong transcriptome-proteome co-evolution along the phylogeny. The top mRNA∼protein co-evolving genes are WDR13, XRCC5, GCN1, and others (**Figure S3c)**. To compare co-evolution between strongly correlated (R>0.8) and non-correlated (|R|<0.2) genes, we determined the signed geometric mean of mRNA and protein PIC values as a proxy, which integrates the direction and size of evolution. We found that highly correlated mRNA-protein pairs tend to co-evolve at all episode nodes throughout the phylogenetical tree (**Figure 2a** and **Figure S4** for C12-C21). For example, at Node C13 which separates EAOG and LAUT clades, geometric means of mRNA and protein PIC are mostly positive and much higher if mRNA∼protein correlation is high (P=2.4e-13) (**Figure 2a**). By mapping the number of protein-protein interactions in STRING (*24*) among proteins, we found that correlated evolution of RNA and protein levels (e.g., R>0.8 vs. |R|<0.2, **Figure 2b**) is prevalent among proteins with a low number of protein-protein interactions.

**Fig. 2.**
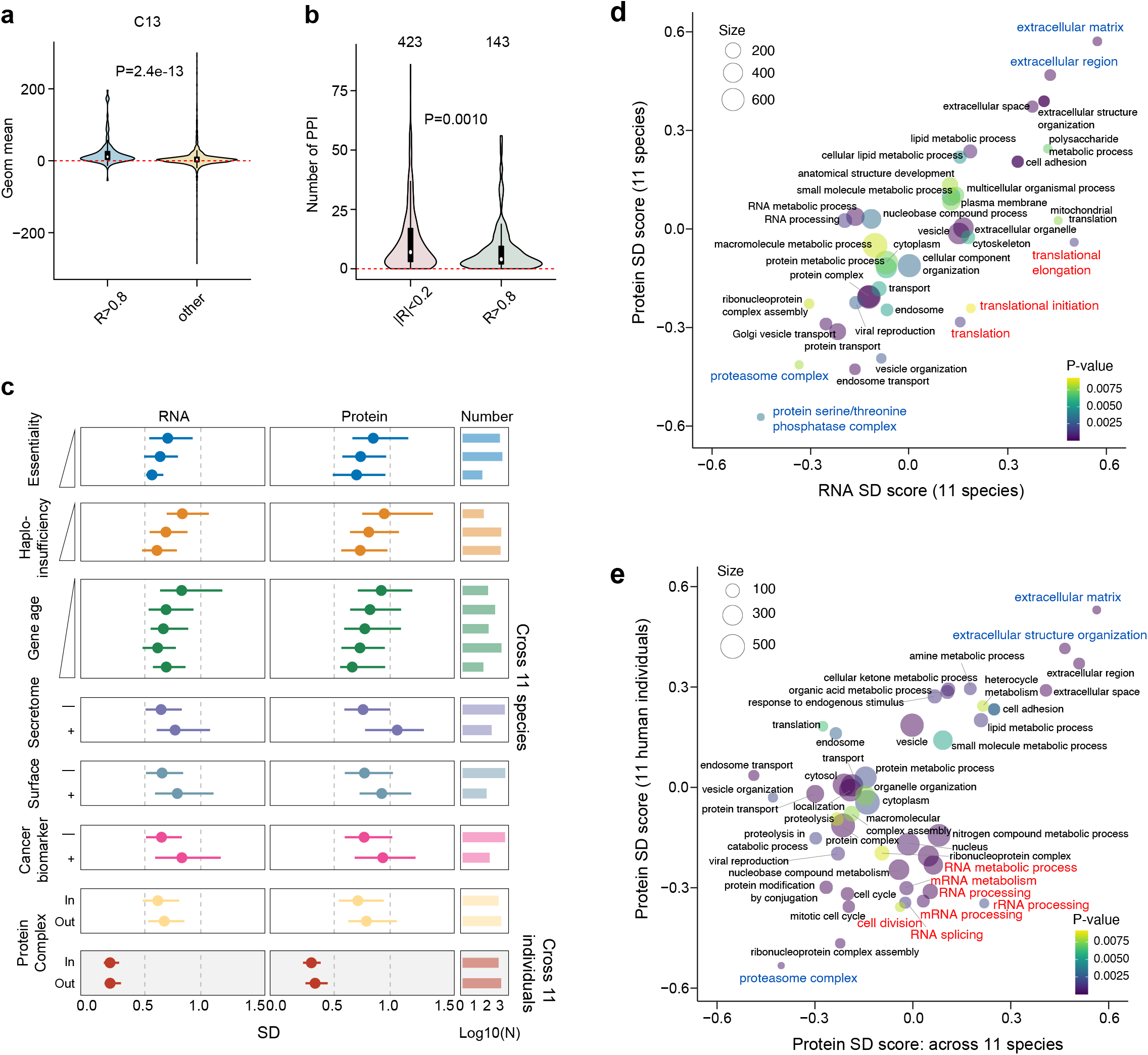
Biological features associated with gene expression and co-evolution in 11 mammalian species at RNA and protein levels. **(a)** Highly correlated mRNA-protein pairs tend to co-evolve. The distribution of signed geometric mean of mRNA-and protein-PIC values at Node C13 for each gene with high mRNA-protein PIC correlations (Pearson’s r > 0.8) and other genes. Node C13 separates Laurasiatheria from Archontoglires. P values, Wilcoxon sum test. **(b)** Number of protein-protein interactions (PPI) of each protein with high (Pearson’s r > 0.8) or low (|Pearson’s r| <0.2) mRNA-protein PIC correlations. P values, Wilcoxon sum test. **(c)** Profiling of SD for mRNA and protein levels and their relationship to gene essentiality (Wilcoxon sum test P = 9.948e-09 and 0.005369 for nonessential vs. Conditional essential at mRNA level and 2.644e-15 and 0.00082 at the protein level), haplo-insufficiency (P=0.0002776, 3.374e-7 and 7.44e-9 for haplo-insensitive vs. medium, insensitive vs. sensitive, and medium vs. sensitive at mRNA level, and 0.005101, 3.24e-05, and 1.675e-07 at the protein level),, secretome (P=1.137e-08 and 2.30e-15 at the mRNA and protein levels), surfaceome (P=0.0006176 and 0.0004607 at the mRNA and protein levels), cancer biomarker annotations (P=1.51e-7 and 0.0001036 at the mRNA and protein levels) and whether the corresponding proteins are in stable protein complex in the CORUM database (Complex_in) or not (Complex_out). Note, P=0.0004556 and 2.393e-6 were obtained at the mRNA and protein levels respectively for comparing Complex_In and Out groups in Cross-species comparison, and P=0.2582 and 3.816e-6 in the Cross-individual comparison). The point range showed the median, 25th and 75th quantile and the histogram showed gene number in each class. **(d)** Two-dimensional (2D) enrichment plot of SD selected GOBPs or GOCCs. The axes denote enrichment scores for SD from the average at mRNA (x-axis) and protein (y-axis) levels across 11 species. **(e)** 2D enrichment plot of SD selected GOBPs or GOCCs denoting gene expression variability analysis across species and human individuals. The axes denote enrichment scores for the protein SD from average across 11 species (x-axis) and 11 individuals (y-axis).

To discern the functional implications of gene expression variability, we calculated the standard deviation (SD) for mRNA and protein levels among species (see **Methods**). The levels of essential or conditionally-essential (*25*) gene products were verified to be much less variable during evolution than non-essential gene expressions (**Figure 2c**, all *P* values in Legend). Likewise, haploinsufficiency means that a single copy of a gene is not sufficient to maintain the normal function. And haploinsufficient genes (*26*) are expressed with much more stable transcript and protein levels. Also, the newly evolved *mammal-specific* genes during evolution (i.e., “younger” genes) (*27*) show higher expression divergence among mammals – especially at the protein level, than those “older” *eukaryote-specific* genes (**Figure 2c and S5a**). Furthermore, secretory (*28*) and cell surface proteins (*29*) display much less stability between species compared to non-secretory or proteins inside the cells, consistent to their adaptable roles. The reported cancer biomarkers (*30*) also showed larger variability, indicating they might be prone to be dys-regulated in disease states. Together, the inter-species SD of mRNA and protein abundances provides an evolutionary angel to understand gene functional diversity.

To broadly map mRNA and protein species-variability to gene functions in an unbiased manner, we utilized a two-dimensional (2D) enrichment plot (*31*) (**Figure 2d**). ***Firstly***, we found that most functional classes exhibit highly-correlated mRNA and protein variabilities – when mRNA levels are variable, the protein concentrations also tend to be diverse across species, and vice versa. This overall intriguing trend again endorses the strong co-evolution of transcriptome-proteome across gene classes (**Figure S5b)** (*2*). ***Secondly***, among the gene classes, extracellular matrix-related have most variable expressions at both transcript and protein levels (highlighted in *blue*, **Figure 2d**). This agrees well with the role of extracellular matrix in interacting to environment. On the other hand, the cellular protein and phosphorylation removal systems such as the proteasome complex and the Serine/Threonine phosphatase complexes showed the least variable concentrations among all functional classes. This may imply that those cell adaptive responses, correcting excessive protein copies (*32*) and abnormal phosphorylation (*33*), are evolutionarily conserved. ***Thirdly***, the most deviated categories from transcriptome-proteome co-evolution are translation-related especially translational elongation and initiation, for which the protein variability is strongly buffered compared to that of mRNA.

In summary, both PIC analysis and functional annotations of co-varying mRNAs and proteins demonstrate a global co-evolutionary dynamics of transcriptome and proteome across gene classes among mammals, with deviating classes mainly due to the protein level buffering and protein-protein interaction constraints.

### Biological variability analysis: Inter-individual versus inter-species

How to understand the evolution-associated gene expression variability, if compared to variation at other scales? We herein referred to a published dataset, part of which reported the proteome variability among 11 unrelated healthy human individuals by using the same cell type and DIA-MS technique (*34*). This comparison thus presents an inter-species (n=11) versus inter-individual (n=11) scrutiny on biodiversity. Combined PCA analysis (see **Methods**) indicates that the global variability between mammalian species is conceivably much larger than the variability between human individuals (**Figure S5c**). Besides the extent of variation, proteins participating in any heteromeric protein complex (i.e., Complex_In, according to the annotation in CORUM (*35*)) exhibited significantly lower inter-individual and inter-species SDs than other “Complex_Out” proteins (**Figure 2c**, bottom two panels). This globally attests the proteostasis control through protein complex stoichiometry, which was previously found post-transcriptionally (*34, 36-38*). However, our data indicates that the mRNA variability in the Complex_In group is also notably lower than that of Complex_Out. The gene-gene correlation analysis demonstrates the consistent trend (**Figure S2d**). Hence, the evolutionarily diversity might have already started to intensively shape the transcript abundance towards protein-level usage of e.g., protein complexes.

Next, we used 2D enrichment plot (*31*) to illustrate the functional convergence between inter-individual diversity and inter-species diversity at the proteome level (**Figure 2e**). We found that, at both individual and species scales, the proteasome complex, again, manifests the lowest protein abundance variability, whereas the extracellular matrix shows the highest. Compared to the same items enriched in **Figure 2d**, it is thus appealing to deduce that abundance control of the subcellular proteomes can extend over multiple scales of biodiversity (**Figure S5d**). Very interestingly, RNA processing and cell division pathways show a particular higher protein abundance variability at the species-scale than individual-scale, indicating their supreme roles during mammalian proteotype evolution. In addition, at the mRNA level, a similar analysis suggests that while the most variable classes are again extracellular matrix related at both scales, the most stable classes are RNA processing-associated pathways (**Figure S6**). Thus, during mammalian evolution, the RNA processing and splicing, although stable between individuals, are tightly regulated at the protein level.

To summarize, the inter-individual and inter-species proteome variabilities are in fact globally correlated and function dependent.

### Diverse evolutionary conservation for proteasome-and lysosome-mediated protein degradation

Due to the prominent steadiness of proteasome expression between individuals and between species, we next sought to interrogate the inter-species stability of the general protein degradation machineries in the cell. We found that the transcript and protein abundance profiles of lysosomal hydrolases and ubiquitin (Ub) enzymes including both deubiquitylating enzymes (DUBs) and E3 Ubiquitin ligases demonstrate distinctly higher variabilities than the proteasome components across individuals and species (**Figure 3a** and **Figure S7**). Whereas the distribution of Ub enzyme levels is wide, the lysosomal proteases display an even higher protein expression diversity than the average level proteome-wide (P=5.38e-05, **Figure S7**). This striking result suggests that the lysosome-mediated degradation is heavily evolving in mammals, despite proteasomal degradation is evolutionarily conserved. Only two proteasome proteins showed exceptional instability – PSMA4 and PSME2, the latter was implicated in immunoproteasome assembly (*39*). Many lysosome hydrolases seem to be expressed lower in Opossum than EAOG and LAUT species in which they become more variable. Certain DUBs such as USP5, 7, 8, 14, and 47 are as conserved as proteosome core subunits, whereas USP 19 and 48 are quite dynamic among mammals. A summary network analysis not only reinforces similar rates of mRNA and protein divergence in these cellular protein removal processes, but additionally reveals that a few Ub enzymes such as VCPIP1, OTULIN, XIAP, and TRIP12 underwent extensive post-transcriptional regulation during evolution (**Figure 3b**).

**Fig. 3.**
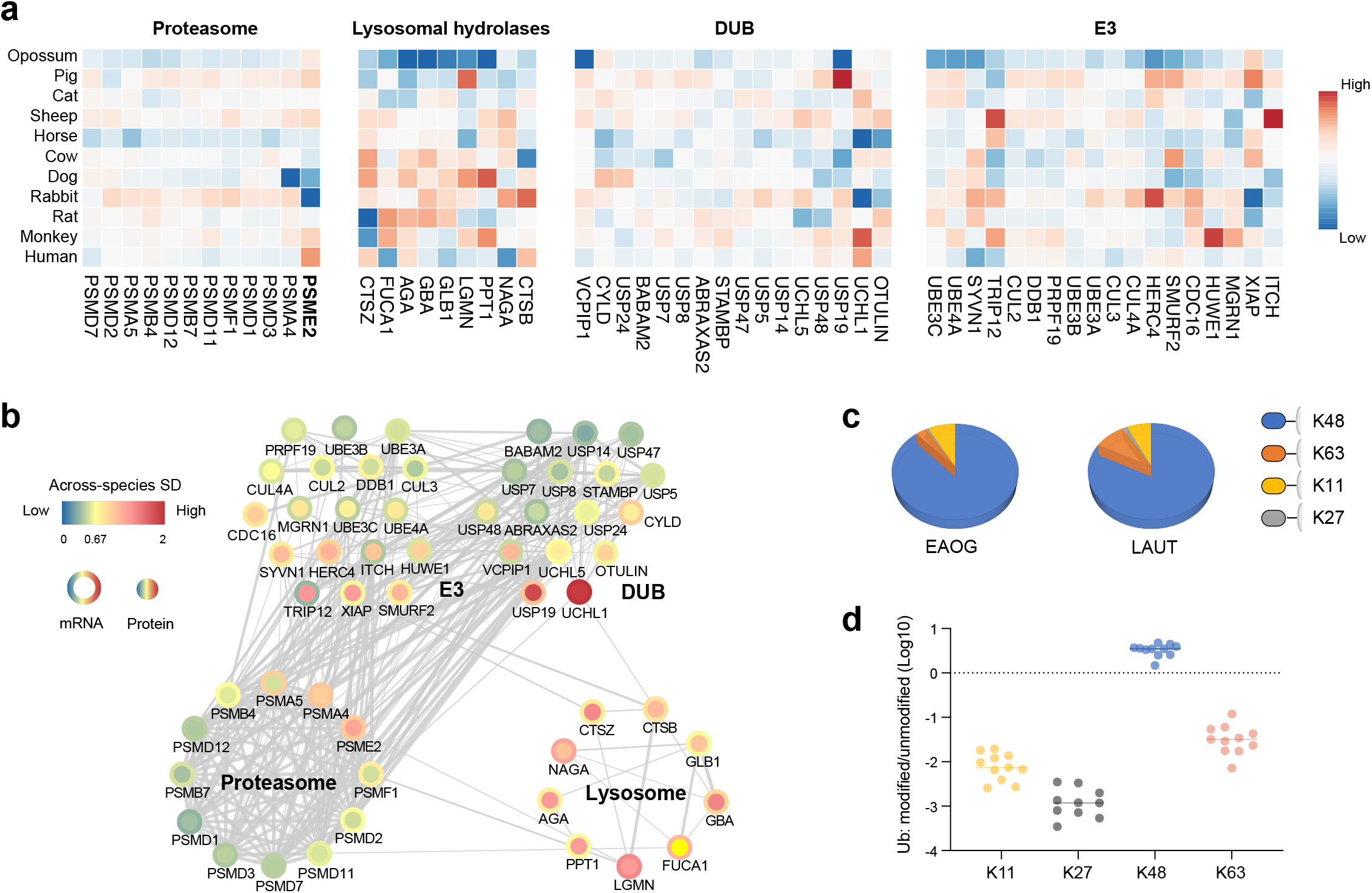
The distinctive inter-species quantitative diversity for proteasomal and lysosomal protein degradation. **(a)** Heatmaps visualizing the protein-specific variability in 11 species for proteins detected and annotated as Proteasome, Lysosomal hydrolases, ubiquitin (Ub) enzymes including both deubiquitylating enzymes (DUBs) and E3 Ubiquitin ligases. **(b)** Network analysis visualizing STRING interactions between proteins. The outer ring and inner circle are colored for mRNA and protein SDs respectively. **(c)** The pie plot visualizing the absolute DIA-MS signals (normalized to total signal per sample) for the signature modified peptides quantifying the Ub chains with specific linkage to different Ks. **(d)** The log-scale ratios of modified Ub peptide vs. unmodified peptide reference for K11, K27, K48, and K63 that were quantified across species.

To corroborate these results, we performed an independent ubiquitin proteomics analysis (*40*), to quantify Ub chains with specific Ub-chain linkage to different Lysine (K) residues. We were able to detect the signature peptides representing Ub modified lysine positions, including K6, K11, K27, K48, and K63 – the later four were successfully quantified among all species with the same modified peptide and unmodified peptide as the relative quantification reference (see **Methods**). We found roughly >10-80 times more K48-linked Ub chains than the other three forms based on DIA-MS signals, strongly indicating the dominant presence of K48-linked Ub chains across mammals. The average abundance of K63-linked seems to be higher in LAUT than EAOG clade (**Figure 3c**). Notably, the K48-linked Ub chain copies also demonstrate the apparently smallest variability among the four types (**Figure 3d**). Due to the canonical role of K48 polyUb chains in proteasome-mediated degradation and the diverse signaling roles of K11, K27, and K63, the Ub proteomics quantification agrees well to the above variability analysis.

### Common and co-varying phosphoproteomic signatures across mammalian species

To interrogate how phosphoproteomics could shed light on molecular evolution, we first evaluated cross-species SDs among transcripts, proteins, and phosphoproteins. We found that P-sites are molecularly much more dynamic than mRNAs and proteins (P<2.2e-16, **Figure 4a**). And the protein abundance “regressed out” P-site values (or, P-site_reg) (*41*) still credibly yield higher SDs (P<2.2e-16), indicating that phosphoproteomes captured activity dynamics that is not reflected by transcript and protein abundances. Following, we focused on the 611 P-sites that were aligned across 11 species (hereafter, common P-sites). Compared to all the other P-sites we detected in Human, the common P-sites are predicted with an overall lower “sift_ala_score”, a computational score predicting the system tolerance if the phosphosite residue is mutated to alanine (*42, 43*) (P=6.6e-12) (**Figure 4b**). Besides, the common P-sites exhibit a noteworthy higher *functional fitness score* (*42*) than other P-sites (P=3.6e-10). Moreover, by mapping to a dataset reporting P-site-specific melting temperature (T_m_) (*44*), we found a small but significant difference, indicating that the common P-sites may bring more structural thermal stability to proteins than other P-sites (P=3.5e-4). These results supported the conservative and essential role of common P-sites in evolution and organism fitness. As for sequence features, the frequency of amino acids (A.A.) surrounding all the common P-sites revealed a diverse A.A. distribution and enriched a few signature motifs **(Figure 4c-d**). These motifs, such as (SP), (SP.R), and (R..S), are closely matching to the substrate motifs of Cyclin-dependent kinases (CDK) and Calmodulin-dependent Protein kinase (CaM) or Protein kinase A (PKA). Lastly, a relative A.A. frequency comparison to human background discovered significant depletion of Glutamic acid (E) and enrichment of Arginine (R) and Serine (S) around the common P-sites (**Figure 4e**). These discovered motifs and rules might be useful for predicting P-site evolutionary conservation in mammals.

**Fig. 4.**
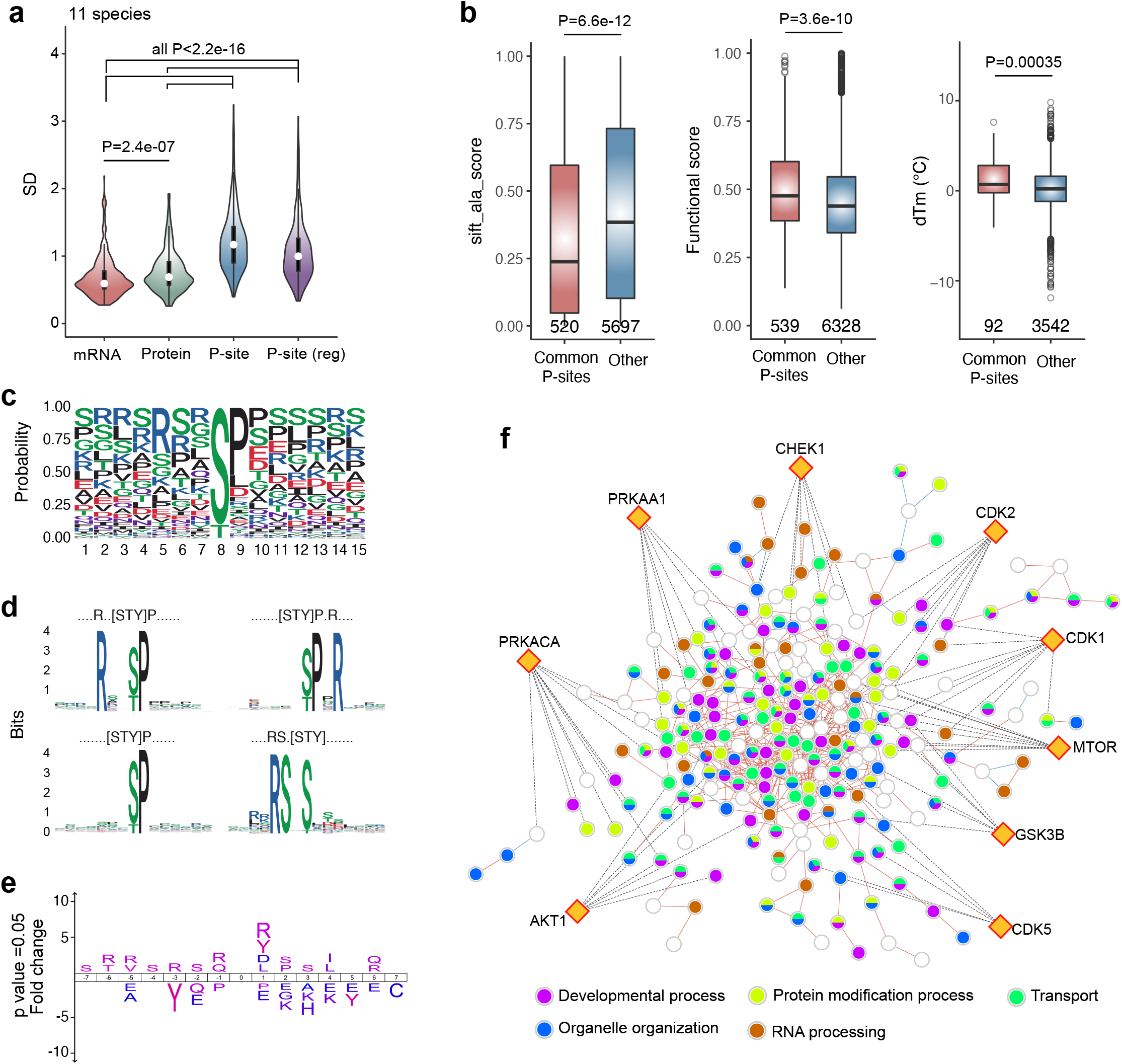
Phosphorylation sites characteristics among 11 mammalian species. **(a)** Profiling of SD across 11 species from the average at mRNA, protein, P-site, and protein-corrected P-site levels from the intersecting list between layers. **(b)** The site-specific parameters and features of the common P-sites (detected in all 11 species) and other human P-sites. P values, Wilcoxon sum test. **(c)** The Sequence logo of surrounding amino acids (±7 aa) of common P-sites (detected in all 11 species). **(d)** Motif analysis of the flanking amino acids (±7 aa) around the common P-sites (detected in all 11 species). The representative enriched motifs (four motif examples with >7 fold of enrichment) were shown. **(e)** Sequence analysis of the flanking amino acids (±7 aa) around the P-sites (common P-sites versus other human P-sites). The fold changes of significant residues (p < 0.05) were shown. **(f)** By inferring the binary, quantitative correlation analysis between any P-site pairs, the phosphorylation co-evolution network was built on the significantly associated common P-sites (Pearson’s correlation, p < 0.001) after correction by total protein changes. Red lines indicate positive associations while blue lines indicate negative associations. The representative GOBPs of phosphoproteins were highlighted in different colors. The top 9 kinases curated from the P-sites as substrates were shown as diamonds and the kinase-substrates pairs were shown as dashed lines.

Previously, little has been known about the structure and evolution of phosphorylation signaling networks. We herein built a phosphorylation co-evolution network (**Figure 4f**, and **Figure S8** for all P-site identities). Each node represents a particular P-site. And each edge denotes a strong *site-to-site* Pearson correlation (P<0.001) based on the P-site_reg levels across the 11 mammalian species. We found the majority (i.e., 96.23%) *site-to-site* correlations co-evolving in the network are strongly positive. Also, the biological annotation suggests that most of the P-sites (i.e., 74.85%) could be functionally annotated to five processes including transport, RNA processing, protein modification process, organelle organization, developmental process. The top nine kinases, including three CDKs (1, 2, and 5), could govern 18.69% of the P-sites in this network (**Figure 4f**), indicating their prevalent roles in mammalian evolution.

In summary, our phosphoproteomics discovered conservative P-sites motifs and a pilot phosphorylation co-evolutionary network containing variance independent of protein abundance.

## Discussion

Our *de-facto* proteomic data for the first time delineate both the protein expression and activity regulations underlying evolutionary diversity. In agreement with Wang et al (*2*), we found that the old, essential, and housekeeping genes tend to express with much smaller between-species divergence than other genes. We also found that co-evolution of expression layers is strong and across gene classes. In our results, the global proteome variability is comparable or even slightly higher than the transcriptome level across species (**Figure 2c and 4a**), emphasizing the evolution of lineage adaptions, rather than the genome-wide compensatory evolution (*2*), at the proteome level. The data further suggests that the involvement in protein complex or interactions essentially presents additional constraints, which may impair the transcriptome-proteome co-evolution.

The biological variabilities between individuals and between species jointly shape the biological diversity in the Earth. Our data essentially addressed two principles underlying biodiversity. ***The first principle*** is that the proteome largely preserved variability preference from the transcriptional level. As shown in **Figure 2d**, gene expression robustness at mRNA and corresponding protein levels is overall tightly controlled within classes of genes of the same molecular functions. The dynamics of gene products participating in the extracellular matrix could render each mammalian species a rapid environment adaptive response. The proteasome homeostasis seems to be evolutionarily conserved as reported (*32*), with the regulation starting from the transcriptional level according to our data, due to the high cost of protein synthesis (*45*). In accordance, the translation-relevant gene expressions are particularly stabilized at the protein level across species (**Figure 2d**). This reinforces the previous finding that translation efficiency profile is highly conserved in evolution (*46*).

In a mammalian cell, the proteasome and lysosome represent the two major machineries for protein degradation (*47*). Previously, ubiquitin (Ub)-proteasome pathway was found to be evolutionarily conserved (*32*), but little is known about the evolutionary conservation of the lysosomal proteolysis pathway and entire Ub system. We discovered a much higher variability of the lysosomal degradation pathway. Considering the endosome transportation-associated protein expressions are quite stable across species, our discovery might be associated with lysosomal exocytosis for remodeling extracellular proteins (*48*). Consistent to this finding, previous studies about the ubiquitin code (*49*) suggested that K48-linked ubiquitin, the mostly stable Ub codes we determined here, is mainly involved in canonical protein degradation via proteasome. In contrast, other Lysine linked Ubs carrying diverse functions, such as endocytosis and DNA-damage responses (K63), cell-cycle regulation (K11), and innate immunity (K27), were quantitively more variable between species. The lower abundance of K63-linked Ub in EAGO species remains to be evaluated in more species. Even for proteasome itself, the immunoproteasome components such as PSME2 (*11*) manifested high inter-species variation. Altogether, our data delineates a complex and manifold relationship between protein degradation and biodiversity.

***The second principle*** underlying biodiversity is that proteome variability universally emerges at both individual and species scales (**Figure 2e**). Although it is expected that inter-species protein variability globally exceeds the inter-individual variability, proteins in extracellular matrix and proteasome again showed highest and lowest variable abundances respectively also among humans. Those classes such as mRNA splicing, mRNA metabolism, and cell division which showed exceptional higher inter-species variability are intriguing and warrant future investigations. For example, Keren et al have summarized various changing models of mRNA alternative splicing (AS) in different eukaryotic lineages (*50*). AS provides a strategy for relaxing negative selection pressure against evolutionary changes (*51*). In our data, the evolutionary variability of AS pathway is strong at the protein level, which might enhance evolvability and proteotype diversity. The cell division-associated protein variability among mammals was further supported by phosphoproteomics data. Our cross-scale analysis represents a critical step toward comprehensive understanding of *inherent protein variability*. This means that, many proteins might be just easier for the cells to regulate using common molecular machineries, and the evolutionary selective forces would prefer to simply increase the magnitude of their protein diversity, such as the extracellular matrix-mediated pathways. The other type of proteins is particularly regulated along the evolution axis, irrespective of their relative, basic biological variability (or robustness), such as RNA processing. It is thus essential to classify two types of protein variabilities for future evolutionary research.

Compared to early studies (*52*), our phosphoproteome data is fairly large in coverage and quantitative. Previously, little was known about the evolution of phosphorylation signaling networks. Our co-variance analysis between P-sites extracted a first evolutionary-associated phosphorylation network independent of protein levels. The enriched motifs from the common P-sites pointed out similar kinases such as CDK1 and PKA heavily evolving in mammalian evolution. We found the patterns of depleting Glutamic acid (E) and enriching Arginine (R) around the common P-site. This result agrees well to our previous study which showed that such patterns tend to accelerate protein turnover when the P-sites get phosphorylated (*53*). Thus, the timely phosphoprotein turnover may render active selection on P-sites in mammalian evolution.

The present study only analyzed fibroblast cells, whereas enormous single-cell and multi-tissue studies have demonstrated the complexity and diversity of gene expression among different cell types (*14-16*). Although most of our conclusions, such as transcriptome-proteome co-evolution and biological variability control, should not be restricted to fibroblast cells, our results promisingly anchor future proteotype characterization studies on other cell types and multiple tissues across species. In addition, certain evolution and co-evolution molecular events might be more apparent with studies measuring the disturbed or dynamic systems. Moreover, as already shown in studies analyzing protein turnover (*9, 54*), thermal stability (*55*), and protein-protein interaction (*56*), the MS-based quantitative proteomic analysis across multiple species will provide additional insights in answering fundamental evolutionary questions and beyond. We expect the establishment of quantitative landscape of proteins and post-translational modifications across species to further contribute to understanding biological variability and biodiversity on Earth.

## Author contributions

Conceptualization: YL, GPW

Methodology: AD, WL, SW, GPW, YL

Investigation: QB, YH, WL, JM, IP

Visualization: QB, YL, GPW

Funding acquisition: YL, GPW

Supervision: YL, GPW

Writing – original draft: QB, YL

Writing – review & editing: JM, GPW

## Supplementary Materials

Figs. S1 to S9

### Materials and Methods

#### Skin fibroblast cell culture

Human skin fibroblast cells (SF) were purchased from ATCC (CRL-4001). Cow (Bos taurus), dog (Canis lupus), horse (Equus caballus), cat (Felis catus), monkey (Mucaca mulatta), opossum (Monodelphis domestica), rabbit (Oryctolagus cuniculus), sheep (Ovis aries), rat (Rattus norvegicus), and pig (Sus scrofa) SFs were obtained from fresh skin tissue. Briefly, a small piece of skin was collected with hair removed, washed in PBS buffer, and cut into strips approximately 1.0 cm^2^. Dermis was separated from epidermis by enzymatic digestion (30 min in 0.25% Trypsin buffer at 37 °C, followed by dissociation buffer (1 mg/mL collagenase, 1 mg/mL Dispase, 400 μg/mL DNase I) for 45 min at 37 °C. Epidermis was then removed and 2 mm pieces were cut from the dermis and transferred to a 12-well plate and covered with media. Fibroblasts emerged from the explants and grew to confluency in growth media with extra tissue removed. Fibroblast cell cultures were then established in 10-cm dishes with in DMEM with high glucose supplemented with 10% FBS at 5% CO_2_. The sample preparation procedures were approved by Institutional Animal Care & Use Committee (IACUC) under the No. 2021-11483. Three replicates (each of 60-80% confluency) were processed for RNA sequencing and proteomics analyses respectively, with the exception that five replicates processed for human and two for monkey proteomic profiling. After estimating the experimental reproducibility (**Figure S1**), all the replicate measurements were accepted and averaged per each of the 11 mammalian species for transcriptomic and proteomic profiling.

#### RNA isolation and sequencing

RNA was isolated using RNeasy micro kit (QIAGEN) and resuspended in 15 μL of water. The RNA samples were measured at The Yale Center for Genome Analysis on the Agilent Bioanalyzer 2100 to determine RNA quality, prepared mRNA libraries and sequenced on Illumina HiSeq2500 to generate 30–40 million reads per sample (Single-end 75 base pair reads).

#### RNA sequencing data procession

RNA-seq data obtained was quantified using the transcript-based quantification approach provided in the ‘kallisto’ program (*57*). Reads are aligned to a reference transcriptome using a fast hashing of k-mers together with a directed de Bruijn graph of the transcriptome. This rapid quantification technique produces transcript-wise abundances which are then normalized and mapped to individual genes and ultimately reported in terms of TPM (*58*). The Ensembl release 100 (May 2020 version) (*59*) gene annotation model was used and raw sequence reads (single end 75 bp) for SFs from the 11 species were aligned to GRCh38.p13, ARS-UCD1.2, CanFam3.1, Felis_catus_9.0, EquCab3.0, Mmul_10, MonDom5 (Release 97), OryCun2.0, Oar_v3.1, Rnor_6.0 and Sscrofa11.1 reference transcriptome assemblies. In order to facilitate gene expression across species, a one-to-one ortholog dataset consisting of 8216 genes was formulated across the 11 species such that the sum of TPMs across these genes for each species totals to 10e6. The TPMs across replicates of the same species were averaged and log2 transformed for following bioinformatic analysis.

#### Protein extraction, alkylation, and digestion

The fibroblast cell proteomes were harvested as previously described (*34*). Cells were washed with PBS twice and scaped off from the dish using the lysis buffer containing 8M urea containing complete protease inhibitor cocktail (Roche) and Halt™ Phosphatase Inhibitor (Thermo). The cell pellets were then ultrasonically lysed at 4 °C for 2 min using a VialTweeter device (Hielscher-Ultrasound Technology) and centrifuged at 18,000 × g for 1 hour to remove the insoluble material.

Protein concentrations were then determined with a Bradford assay (Bio-Rad, Hercules, CA, USA). The supernatant protein samples were reduced with 10 mM Dithiothreitol (DTT) for 1 h at 57 °C and alkylated by 20 mM iodoacetamide in the dark for 1 h at room temperature. All samples were further diluted by 5 times using 100 mM NH4HCO3 and were digested in-solution with sequencing-grade porcine trypsin (Promega) overnight at 37 °C as described (*36*). The resulted peptide mixture was desalted with a C18 column (MarocoSpin Columns, NEST Group INC). The final peptide amounts were determined by Nanodrop (Thermo Scientific).

#### Phosphopeptide enrichment

Besides ∼4μg of peptides digested per sample that were used for proteomic analysis, all the peptides from different replicates of equal amount were pooled for phosphopeptide enrichment and phosphoproteomics, due to the limited peptide amounts yielded in individual replicate. The phosphopeptide enrichment was performed using the High-Select™ Fe-NTA kit (Thermo Scientific, A32992) according to the manufacturer’s instructions (*60*). Briefly, the resins of spin-column in the Fe-NTA kit were aliquoted and incubated with 80-300 µg of total peptides for 30 min at room temperature. The resins were then transferred into a filter tip (TF-20-L-R-S, Axygen), so that the supernatant was removed by centrifugation. Then, the resins were washed sequentially with 200 µL of washing buffer (80% ACN, 0.1% TFA) by 3 times and 200 µL of LC-MS grade H_2_O by 2 times to remove the nonspecifically adsorbed peptides. The enriched phosphopeptides were then eluted off the resins by 100 µL of elution buffer (50% ACN, 5% NH_3_•H_2_O) 2 times. All the centrifugation steps were kept at 500 g, 30 sec. The eluates per species were combined and dried by speed-vac and stored in −80 °C.

#### Liquid chromatography (LC) settings before mass spectrometry

All peptide-level samples (and their pooled mixtures per each species) were resolved in 2% ACN, 0.1% FA for LC-MS measurements, with 2 µg of peptides or 0.2-0.5 µg of the enriched phosphopeptides injected per measurement. The single-shot DIA-MS analysis of 2.5 hours was performed as previously described (*4, 61*). The LC used was an EASY-nLC 1200 system (Thermo Scientific, San Jose, CA) harboring a 75 µm × 50 cm C18 column packed with 100A C18 material. A 150-min LC separation was configured based on the mix of buffer A (0.1% formic acid in H_2_O) and buffer B (80% acetonitrile containing 0.1% formic acid): Buffer B was made to increase from 4% to 34% in 139 mins, then to surge to 100% in 3 mins, and then kept at 100% for 8 mins. The LC-MS flow rate was kept at 300 nL/min with the temperature-controlled at 60 °C by a PRSO-V1 column oven (Sonation GmbH, Biberach, Germany). The additional column re-equilibrating was performed in about 10-15 mins using the high-flow rate up to ∼800 nL/ min.

#### DIA-MS measurements

The Orbitrap Fusion Lumos Tribrid mass spectrometer (Thermo Scientific) instrument coupled to a nanoelectrospray ion source (NanoFlex, Thermo Scientific) was used as the DIA-MS platform for both proteomic and phosphoproteomic analyses (*4*). Spray voltage was set to 2,000 V and heating capillary temperature at 275 °C. All the DIA-MS methods consisted of one MS1 scan and 33 MS2 scans of variable windows by quadrupole isolation (*62*). This schema was comprised of 350∼373.775, 373.25∼393.75, 393.25∼410.75, 410.25∼427.75, 427.25∼443.75, 443.25∼459.75, 459.25∼474.75, 474.25∼489.75, 489.25∼503.75, 503.25∼518.75, 518.25∼533.75, 533.25∼547.75, 547.25∼562.75, 562.25∼577.75, 577.25∼592.75, 592.25∼608.75, 608.25∼623.75, 623.25∼639.75, 639.25∼656.75, 656.25∼674.75, 674.25∼692.75, 692.25∼711.75, 711.25∼732.75, 732.25∼754.75, 754.25∼778.75, 778.25∼803.75, 803.25∼833.75, 833.25∼866.75, 866.25∼905.75, 905.25∼955.75, 955.25∼1023.75, 1023.25∼1134.75, 1134.225∼1065 with 0.5 m/z overlapping between windows. The MS1 scan range was 350 – 1650 m/z and the MS1 resolution was 120,000 at m/z 200. The MS1 full scan AGC target value was set to be 2.0E6 and the maximum injection time was 50 ms. The MS2 scan range was set to be 200 – 1800 m/z and MS2 resolution was 30,000 at m/z 200. The normalized HCD collision energy was set at 28%. The MS2 AGC was set to be 1.5E6 and the maximum injection time was 52 ms. The default peptide charge state was set to 2.

#### Database search for proteomics and phosphoproteomics

DIA-MS data procession were performed using Spectronaut v14 (*17, 63*) with the “DirectDIA”, an optimal spectral library-free pipeline (*64*). For both proteomics and phosphoproteomics results (*4*), the DIA-MS raw data sets were searched directly against the Ensembl species-specific protein fasta files (zipped files with name ending as “pep.all.fa.gz” at useast.ensembl.org/index.html). These files include:

Bos_taurus.ARS-UCD1.2.pep.all.fa (for “cow”)

Canis_lupus_familiaris.CanFam3.1.pep.all.fa (for “dog”)

Cavia_porcellus.Cavpor3.0.pep.all.fa (for “opossum”)

Equus_caballus.EquCab3.0.pep.all.fa (for “horse”)

Felis_catus.Felis_catus_9.0.pep.all.fa (for “cat”)

Homo_sapiens.GRCh38.pep.all.fa (for “human”)

Macaca_mulatta.Mmul_10.pep.all.fa (for “monkey”)

Oryctolagus_cuniculus.OryCun2.0.pep.all.fa (for “rabbit”)

Ovis_aries.Oar_v3.1.pep.all.fa (for “sheep”)

Rattus_norvegicus.Rnor_6.0.pep.all.fa (for “rat”)

Sus_scrofa.Sscrofa11.1.pep.all.fa (for “pig”)

Particularly, for the total proteomic identification in each species, the possibilities of Oxidation at methionine and Acetylation at the protein N-terminals were set as variable modifications, whereas Carbamidomethylation at cysteine was set as a fixed modification. For the phosphoproteomic identification, the additional possibility of Phosphorylation at serine/threonine/tyrosine (S/T/Y) was enabled as the variable modification. For both proteomic and phosphoproteomic datasets per species, both peptide-and protein-FDR (based on Qvalue) were both controlled at 1%. In particular, the PTM localization option in Spectronaut v^14^ was enabled to locate phosphorylation sites (*6, 65*) in each species with the probability score cutoff >0.75(*65*), which ensures only Class-I peptides (*66*), in which each phosphosite is confidently localized onto one S, T, or Y in the peptide sequence, to be identified, quantified and reported in the results. For each localized phosphosite, the corresponding phosphopeptide precursors (if more than one) were averaged for quantification.

#### Database search for Ubiquitin linkage types

In addition, to search and determine the different type of ubiquitin chains via its lysine residue (i.e., the “ubiquitin code’), a “Gly-Gly” or diGly modification was set up as a variable modification in a separated search in all mammalian species, by searching the total proteomic data against the same fasta files. After the identical FDR control (i.e. 1%) at both peptide and protein level, the diGly localization scoring in peptide sequence, the search results were manually inspected. The quantities of K11-, K27-, K48-, and K63-were inferred based on most abundant peptide precursors for TLTGK_GG_TITLEVEPSDTIENVK (K11), TITLEVEPSDTIENVK_GG_AK (K27), LIFAGK_GG_QLEDGR (K48), and TLSDYNIQK_GG_ESTLHLVLR (K63), respectively. To accurately determine the relative quantitative variability between K11-, K27-, K48-, and K63-linked chains, the above peptides carrying diGly were compared to the unmodified counterpart peptide with no miss-cleavage. This means, TLSDYNIQK and ESTLHLVLR were used for K48 and K63, whereas TITLEVEPSDTIENVK was used for K11 and K27 as counterpart peptides.

#### Quantitative proteotype analysis

All the other Spectronaut settings for identification and quantification were kept as default (*4*). This meaning that for example, the “Missed cleavages” allowed was set at 2, the “Inference Correction” was enabled, the “Global Normalization” (on “Median”) was used, the quantification was performed at the MS2 level using peak areas, the “Protein Inference algorithm” was implemented using “IDPicker”, and the Top 3 peptide precursors (“Min: 1 and Max: 3”) were summed based on MS2-level peak areas for representing protein quantities in all DIA analyses. The quantitative data reported by Spectraonuat analysis for proteins and phosphosites were then log2-transformed for downstream statistical analysis if applicable. As for multi-species analysis, due to the database searching against Ensembl species-specific protein fasta files, the proteomic quantitative results could be directly added to the “one-to-one ortholog” data table consisting of 8216 genes (see above for RNA-seq data procession) using the ensemble gene identities, allowing the transcriptomic and proteomic quantifications to be summarized with a “gene-centric” perspective. For the absolute-scale analysis, the log2-transformed mRNA, protein and phosphosite quantification data are compared between molecular layers or between species. For the relative-scale analysis, the mRNA, protein and phosphosite quantification data of each species was compared to the *averaged* values across all 11 species, summarized as fold-changes (FCs) and the log2 transformed FCs (i.e., individual species /averaged values of 11 species) were used for relative correlation analysis and variability analysis (i.e., by determining the Standard deviation, SD).

#### Bioinformatic analysis

Circos-0.69-9 (http://circos.ca) (*67*) was used for the circle visualization (Figure 1a). Functional annotation was carried out in David Functional Annotation Tool v6.8 (https://david.ncifcrf.gov/summary.jsp) (*68*) with all detected proteins in this study as background (Figure S2). Pearson and Spearman’s correlation coefficients were calculated using R (functions cor() or cor.test() to infer statistical significance). Principal component analysis (PCA) was performed by NIA Array Analysis (*69*). The annotation of surfaceome, secretome, cancer biomarkers, gene essentiality, haplo-insufficiency, and age were annotated respectively by Cell Surface Protein Altas (*29*), human secretome map (*28*), Human Protein Atlas (https://www.proteinatlas.org) (*30*), OGEE v2 (*25*), HIPred scores (*26*), and modeAge (*27*) (Figure 2). Protein complex information was extracted from the CORUM database (*35*) (Figure 2). The gene ontology (GO) and Kyoto Encyclopedia of Genes and Genomes (KEGG) annotation and 2D enrichment analysis were performed in Perseus v1.6.8.0 (*70*). The retrieve of flanking amino acid sequences (±7 amino acids) of phosphorylation sites and the motif enrichment were performed by motifeR (https://www.omicsolution.org/wukong/motifeR) (*71*) (Figure 4). Sequence analysis (Figures 4) was conducted and visualized by IceLogo (https://iomics.ugent.be/icelogoserver) (*72*). The functional score can reflect the importance of phosphosite for organismal fitness (*73*) (Figure 4). The sift score predicts the functional impact of missense variants based on sequence homology and the physicochemical properties of the amino acids (*74*) (Figure 4). The melting temperature (Tm, °C) and T_1/2_ value (Figure 4) for each phosphosite was taken from the reported datasets (*44, 53*). The net phosphorylation changes were detected by total protein changes correction through linear regression as reported previously (*41*). After regression, the phospho-site pairs with significant correlation (Pearson, p < 0.001) were used to construct interaction network by Cytoscape (*75*) (Figure 4). The kinase-substrate relations were curated from PhosphoSitePlus (*76*). Gene homology were retrieved from Ensembl (https://www.ensembl.org). Only “one-to-one” orthologs were considered. To identified the homologous phosphor-sites across 11 species from our phosphorylation datasets, the sequence windows (±7aa flanking sequence) of detected phosphor-sites in orthologous proteins were aligned by the “pairwiseAlignment” function in R package “Biostrings”. The sequence windows with score ≥10 and in orthologous proteins were considered as homologous. Data from other species were first mapped to human data, and then the common phospho-sites were obtained as the intersection of all species. Phylogenetically independent contrast (PIC) were computed by R-package APE (Analysis of Phylogenesis and Evolution) using the function “pic” (*77*). The number and relationship of protein-protein interactions (PPI) were curated using STRING v11.0 (https://string-db.org) (*24*).

#### Data availability

Raw data will be available upon paper publication.

#### A website inventory for proteome-centric multi-species navigation

Due to the well-matched multi-omics layers especially the application of consistent DIA-MS, we consider our data set a high-quality resource for future mammalian evolution and gene expression studies. We thus generated a website to facilitate the navigation of the data basis (**Figure S9**). This website interactively provides queries about the abundances for any transcript or protein in every species. It additionally offers heatmaps and scatterplots between molecular layers for individual gene or gene sets of interest.

## References

1. J. B. Muller et al., The proteome landscape of the kingdoms of life. Nature 582, 592–596 (2020).

2. Z. Y. Wang et al., Transcriptome and translatome co-evolution in mammals. Nature 588, 642–647 (2020).

3. R. Aebersold, M. Mann, Mass-spectrometric exploration of proteome structure and function. Nature 537, 347–355 (2016).

4. E. Gao et al., Data-independent acquisition-based proteome and phosphoproteome profiling across six melanoma cell lines reveals determinants of proteotypes. Mol Omics 17, 413–425 (2021).

5. P. Navarro et al., A multicenter study benchmarks software tools for label-free proteome quantification. Nature biotechnology 34, 1130–1136 (2016).

6. G. Rosenberger et al., Inference and quantification of peptidoforms in large sample cohorts by SWATH-MS. Nature biotechnology 35, 781–788 (2017).

7. B. C. Collins et al., Multi-laboratory assessment of reproducibility, qualitative and quantitative performance of SWATH-mass spectrometry. Nature communications 8, 291 (2017).

8. Y. Xuan et al., Standardization and harmonization of distributed multi-center proteotype analysis supporting precision medicine studies. Nature communications 11, 5248 (2020).

9. K. Swovick et al., Cross-species Comparison of Proteome Turnover Kinetics. Molecular & cellular proteomics : MCP 17, 580–591 (2018).

10. P. Picotti et al., A complete mass-spectrometric map of the yeast proteome applied to quantitative trait analysis. Nature 494, 266–270 (2013).

11. N. Romanov et al., Disentangling Genetic and Environmental Effects on the Proteotypes of Individuals. Cell 177, 1308–1318 e1310 (2019).

12. J. D. Venable, M. Q. Dong, J. Wohlschlegel, A. Dillin, J. R. Yates, Automated approach for quantitative analysis of complex peptide mixtures from tandem mass spectra. Nature methods 1, 39–45 (2004).

13. L. C. Gillet et al., Targeted data extraction of the MS/MS spectra generated by data-independent acquisition: a new concept for consistent and accurate proteome analysis. Molecular & cellular proteomics : MCP 11, O111 016717 (2012).

14. C. Ding et al., A Cell-type-resolved Liver Proteome. Molecular & cellular proteomics : MCP 15, 3190–3202 (2016).

15. D. P. Nusinow et al., Quantitative Proteomics of the Cancer Cell Line Encyclopedia. Cell 180, 387–402 e316 (2020).

16. S. Kim-Hellmuth et al., Cell type-specific genetic regulation of gene expression across human tissues. Science 369, (2020).

17. R. Bruderer et al., Optimization of Experimental Parameters in Data-Independent Mass Spectrometry Significantly Increases Depth and Reproducibility of Results. Molecular & cellular proteomics : MCP 16, 2296–2309 (2017).

18. G. Rosenberger et al., Statistical control of peptide and protein error rates in large-scale targeted data-independent acquisition analyses. Nature methods 14, 921–927 (2017).

19. K. L. Howe et al., Ensembl 2021. Nucleic Acids Res 49, D884–D891 (2021).

20. Y. Liu, A. Beyer, R. Aebersold, On the Dependency of Cellular Protein Levels on mRNA Abundance. Cell 165, 535–550 (2016).

21. C. Buccitelli, M. Selbach, mRNAs, proteins and the emerging principles of gene expression control. Nature reviews. Genetics 21, 630–644 (2020).

22. C. Vogel, E. M. Marcotte, Insights into the regulation of protein abundance from proteomic and transcriptomic analyses. Nature reviews. Genetics 13, 227–232 (2012).

23. T. Garland, Jr., A. R. Ives, Using the Past to Predict the Present: Confidence Intervals for Regression Equations in Phylogenetic Comparative Methods. Am Nat 155, 346–364 (2000).

24. D. Szklarczyk et al., STRING v11: protein-protein association networks with increased coverage, supporting functional discovery in genome-wide experimental datasets. Nucleic Acids Res 47, D607–D613 (2019).

25. W. H. Chen, G. Lu, X. Chen, X. M. Zhao, P. Bork, OGEE v2: an update of the online gene essentiality database with special focus on differentially essential genes in human cancer cell lines. Nucleic Acids Res 45, D940–D944 (2017).

26. H. A. Shihab, M. F. Rogers, C. Campbell, T. R. Gaunt, HIPred: an integrative approach to predicting haploinsufficient genes. Bioinformatics 33, 1751–1757 (2017).

27. B. J. Liebeskind, C. D. McWhite, E. M. Marcotte, Towards Consensus Gene Ages. Genome biology and evolution 8, 1812–1823 (2016).

28. M. Uhlen et al., The human secretome. Science signaling 12, (2019).

29. D. Bausch-Fluck et al., The in silico human surfaceome. Proceedings of the National Academy of Sciences of the United States of America 115, E10988–E10997 (2018).

30. M. Uhlen et al., Proteomics. Tissue-based map of the human proteome. Science 347, 1260419 (2015).

31. J. Cox, M. Mann, 1D and 2D annotation enrichment: a statistical method integrating quantitative proteomics with complementary high-throughput data. BMC bioinformatics 13 Suppl 16, S12 (2012).

32. A. Rousseau, A. Bertolotti, An evolutionarily conserved pathway controls proteasome homeostasis. Nature 536, 184–189 (2016).

33. C. Gingras et al., A novel, evolutionarily conserved protein phosphatase complex involved in cisplatin sensitivity. Molecular & cellular proteomics : MCP 4, 1725–1740 (2005).

34. Y. Liu et al., Systematic proteome and proteostasis profiling in human Trisomy 21 fibroblast cells. Nature communications 8, 1212 (2017).

35. M. Giurgiu et al., CORUM: the comprehensive resource of mammalian protein complexes-2019. Nucleic Acids Res 47, D559–D563 (2019).

36. Y. Liu et al., Multi-omic measurements of heterogeneity in HeLa cells across laboratories. Nature biotechnology 37, 314–322 (2019).

37. N. Dephoure et al., Quantitative proteomic analysis reveals posttranslational responses to aneuploidy in yeast. eLife 3, e03023 (2014).

38. C. Juschke et al., Transcriptome and proteome quantification of a tumor model provides novel insights into post-transcriptional gene regulation. Genome Biol 14, r133 (2013).

39. S. Murata et al., Immunoproteasome assembly and antigen presentation in mice lacking both PA28alpha and PA28beta. EMBO J 20, 5898–5907 (2001).

40. W. Kim et al., Systematic and quantitative assessment of the ubiquitin-modified proteome. Molecular cell 44, 325–340 (2011).

41. T. I. Roumeliotis et al., Genomic Determinants of Protein Abundance Variation in Colorectal Cancer Cells. Cell reports 20, 2201–2214 (2017).

42. D. Ochoa et al., The functional landscape of the human phosphoproteome. Nature biotechnology, (2019).

43. P. C. Ng, S. Henikoff, SIFT: Predicting amino acid changes that affect protein function. Nucleic Acids Res 31, 3812–3814 (2003).

44. J. X. Huang et al., High throughput discovery of functional protein modifications by Hotspot Thermal Profiling. Nature methods 16, 894–901 (2019).

45. J. R. Warner, The economics of ribosome biosynthesis in yeast. Trends Biochem Sci 24, 437–440 (1999).

46. T. Tuller et al., An evolutionarily conserved mechanism for controlling the efficiency of protein translation. Cell 141, 344–354 (2010).

47. I. Dikic, Proteasomal and Autophagic Degradation Systems. Annu Rev Biochem 86, 193–224 (2017).

48. B. Tancini et al., Lysosomal Exocytosis: The Extracellular Role of an Intracellular Organelle. Membranes (Basel) 10, (2020).

49. M. Tracz, W. Bialek, Beyond K48 and K63: non-canonical protein ubiquitination. Cell Mol Biol Lett 26, 1 (2021).

50. H. Keren, G. Lev-Maor, G. Ast, Alternative splicing and evolution: diversification, exon definition and function. Nature reviews. Genetics 11, 345–355 (2010).

51. Y. Xing, C. Lee, Alternative splicing and RNA selection pressure--evolutionary consequences for eukaryotic genomes. Nature reviews. Genetics 7, 499–509 (2006).

52. J. Boekhorst, B. van Breukelen, A. Heck, Jr., B. Snel, Comparative phosphoproteomics reveals evolutionary and functional conservation of phosphorylation across eukaryotes. Genome Biol 9, R144 (2008).

53. C. Wu et al., Global and Site-Specific Effect of Phosphorylation on Protein Turnover. Developmental cell 56, 111–124 e116 (2021).

54. K. Swovick et al., Interspecies Differences in Proteome Turnover Kinetics Are Correlated With Life Spans and Energetic Demands. Molecular & cellular proteomics : MCP 20, 100041 (2021).

55. A. Jarzab et al., Meltome atlas-thermal proteome stability across the tree of life. Nature methods 17, 495–503 (2020).

56. C. Wan et al., Panorama of ancient metazoan macromolecular complexes. Nature 525, 339–344 (2015).

57. N. L. Bray, H. Pimentel, P. Melsted, L. Pachter, Near-optimal probabilistic RNA-seq quantification. Nat Biotechnol 34, 525–527 (2016).

58. G. P. Wagner, K. Kin, V. J. Lynch, Measurement of mRNA abundance using RNA-seq data: RPKM measure is inconsistent among samples. Theory in biosciences = Theorie in den Biowissenschaften 131, 281–285 (2012).

59. A. D. Yates et al., Ensembl 2020. Nucleic Acids Res 48, D682–D688 (2020).

60. Q. Gao et al., Integrated Proteogenomic Characterization of HBV-Related Hepatocellular Carcinoma. Cell 179, 561–577 e522 (2019).

61. M. Mehnert, W. Li, C. Wu, B. Salovska, Y. Liu, Combining Rapid Data Independent Acquisition and CRISPR Gene Deletion for Studying Potential Protein Functions: A Case of HMGN1. Proteomics, e1800438 (2019).

62. B. Salovska, W. Li, Y. Di, Y. Liu, BoxCarmax: A High-Selectivity Data-Independent Acquisition Mass Spectrometry Method for the Analysis of Protein Turnover and Complex Samples. Analytical chemistry 93, 3103–3111 (2021).

63. R. Bruderer et al., Extending the limits of quantitative proteome profiling with data-independent acquisition and application to acetaminophen-treated three-dimensional liver microtissues. Molecular & cellular proteomics : MCP 14, 1400–1410 (2015).

64. C. C. Tsou et al., DIA-Umpire: comprehensive computational framework for data-independent acquisition proteomics. Nature methods 12, 258-264, 257 p following 264 (2015).

65. D. B. Bekker-Jensen et al., Rapid and site-specific deep phosphoproteome profiling by data-independent acquisition without the need for spectral libraries. Nature communications 11, 787 (2020).

66. J. V. Olsen et al., Global, in vivo, and site-specific phosphorylation dynamics in signaling networks. Cell 127, 635–648 (2006).

67. M. Krzywinski et al., Circos: an information aesthetic for comparative genomics. Genome research 19, 1639–1645 (2009).

68. W. Huang da, B. T. Sherman, R. A. Lempicki, Systematic and integrative analysis of large gene lists using DAVID bioinformatics resources. Nature protocols 4, 44–57 (2009).

69. A. A. Sharov, D. B. Dudekula, M. S. Ko, A web-based tool for principal component and significance analysis of microarray data. Bioinformatics 21, 2548–2549 (2005).

70. S. Tyanova et al., The Perseus computational platform for comprehensive analysis of (prote)omics data. Nature methods 13, 731–740 (2016).

71. S. Wang et al., motifeR: An Integrated Web Software for Identification and Visualization of Protein Posttranslational Modification Motifs. Proteomics 19, e1900245 (2019).

72. N. Colaert, K. Helsens, L. Martens, J. Vandekerckhove, K. Gevaert, Improved visualization of protein consensus sequences by iceLogo. Nature methods 6, 786–787 (2009).

73. D. Ochoa et al., The functional landscape of the human phosphoproteome. Nature biotechnology 38, 365–373 (2020).

74. R. Vaser, S. Adusumalli, S. N. Leng, M. Sikic, P. C. Ng, SIFT missense predictions for genomes. Nature protocols 11, 1–9 (2016).

75. P. Shannon et al., Cytoscape: a software environment for integrated models of biomolecular interaction networks. Genome research 13, 2498–2504 (2003).

76. P. V. Hornbeck et al., PhosphoSitePlus, 2014: mutations, PTMs and recalibrations. Nucleic acids research 43, D512–520 (2015).

77. E. Paradis, Analysis of phylogenetics and evolution with R. Use R! (Springer, New York, 2006), pp. xii, 211 p.

78. Y. Perez-Riverol et al., The PRIDE database and related tools and resources in 2019: improving support for quantification data. Nucleic Acids Res 47, D442–D450 (2019).

