## Supplementary Figures S1-S9 for "Proteotype Co-evolution and Diversity in Mammals"

Supplementary Figure 1

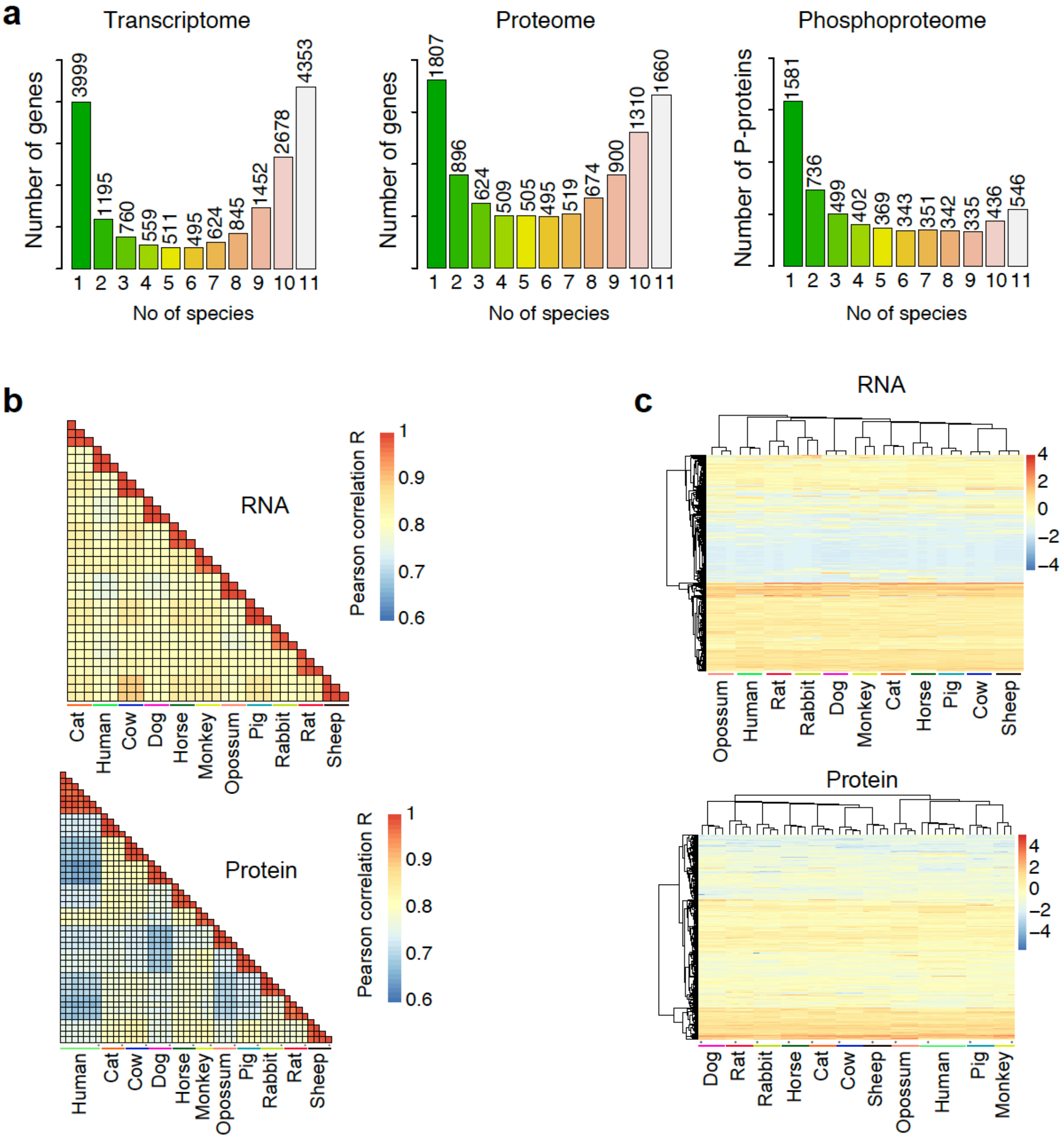


**Figure S1. Overview of transcriptome, proteome and phosphoproteome in 11 mammalian species.**

(a) Identification numbers of mRNA (TPM>1), protein or phosphoprotein detected in single or multiple species (b) The correlation analysis of the RNA and protein profiles among 11 mammalian species. (c) Hierarchical clustering analysis of the RNA and protein profiles among 11 mammalian species.

Supplementary Figure 2

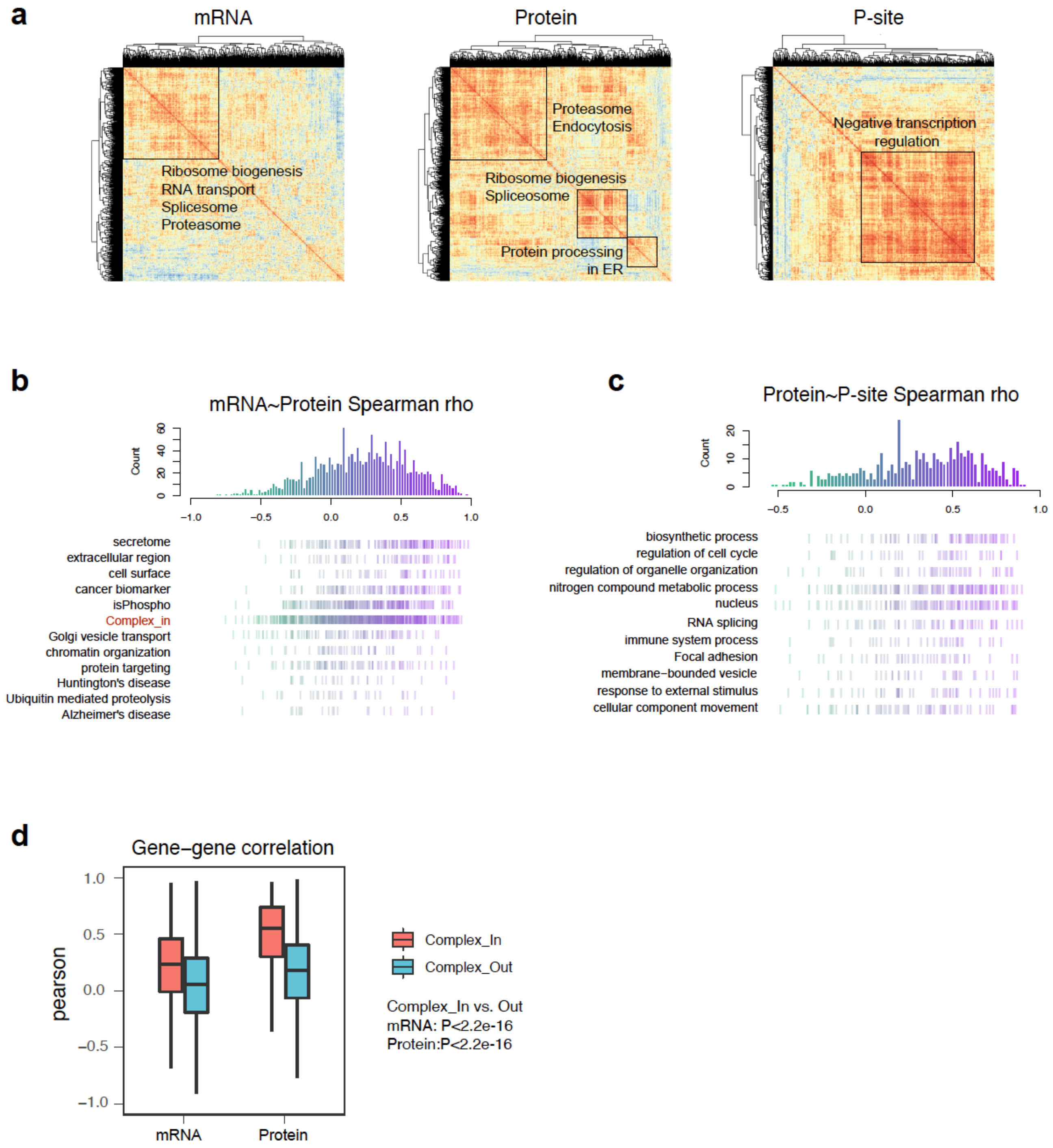


**Figure S2. Relationship between genes and between molecular layers reveals functional diversity.**

(a) Heatmap visualization of covariation among the common genes in 11 species, including 1660 genes detected at both mRNA and protein levels in all species and 611 P-sites detected in all species. Boxes display the highly associated clusters and the key enriched processes. Items of size >15 and $P<0.05$ are shown. (b-c) Representative pathways showing different distribution of mRNA~protein (b) and Protein~P-site (c) correlation. (d) Profiling of gene-gene correlations at mRNA and protein levels and their relationship to Corum Complex (i.e., annotated into a protein complex, Complex_In, or not annotated in any complexes, Complex_Out).

Supplementary Figure 3

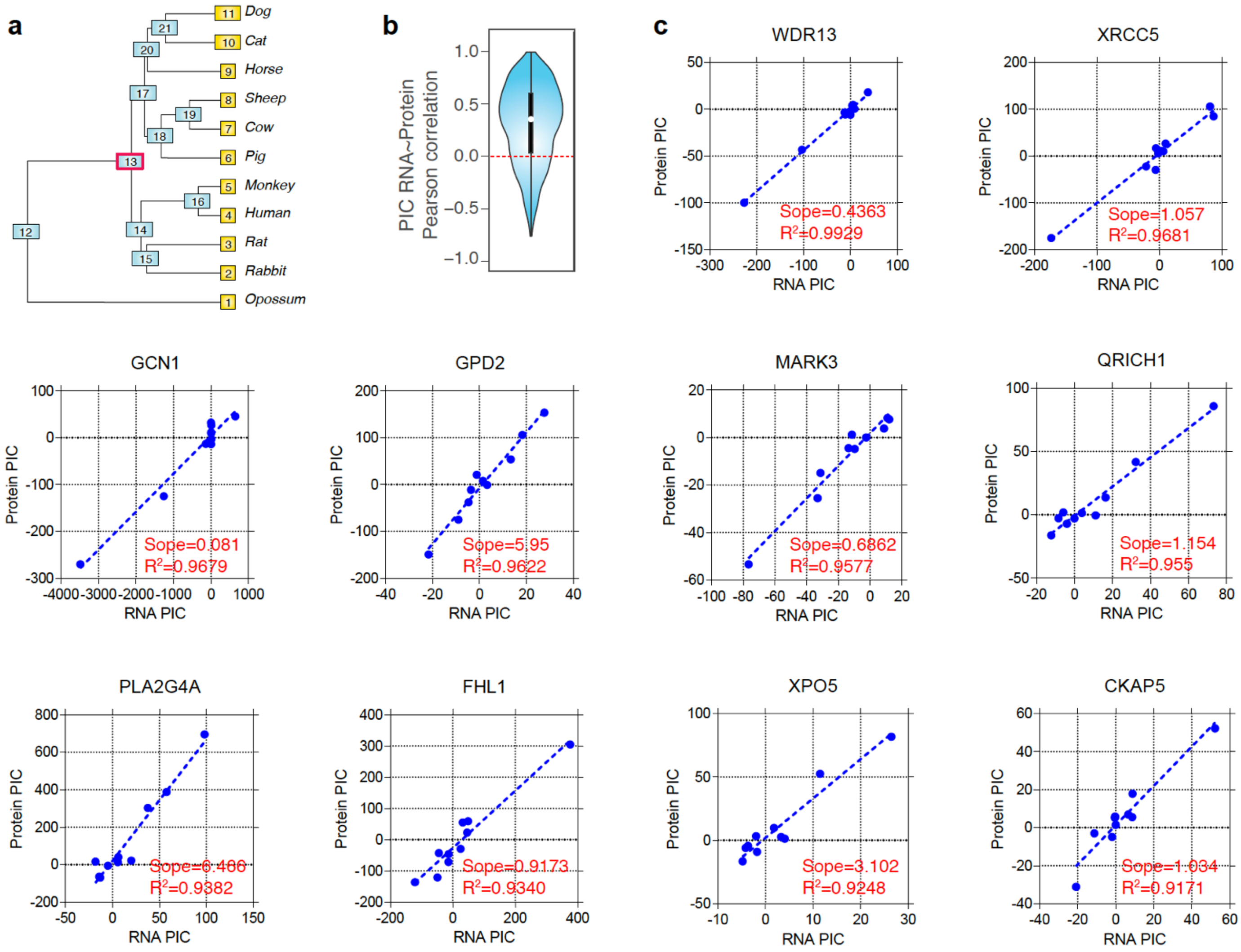


**Figure S3. Phylogenetically independent contrast (PIC) analysis suggests mRNA and proteins are overall co-evolving in 11 mammalian species.**

(a) All the nodes depicting evolutionary relationship between the 11 mammalian species based on the phylogeny tree. For example, node C13 separates Laurasiatheria from Archontoglires in the phylogeny tree. (b) The distribution of Pearson correlation among the RNA-PIC and Protein-PIC of all common genes in 11 species. (c) PIC regression of the top 10 most highly correlated RNA-protein pairs.

Supplementary Figure 4

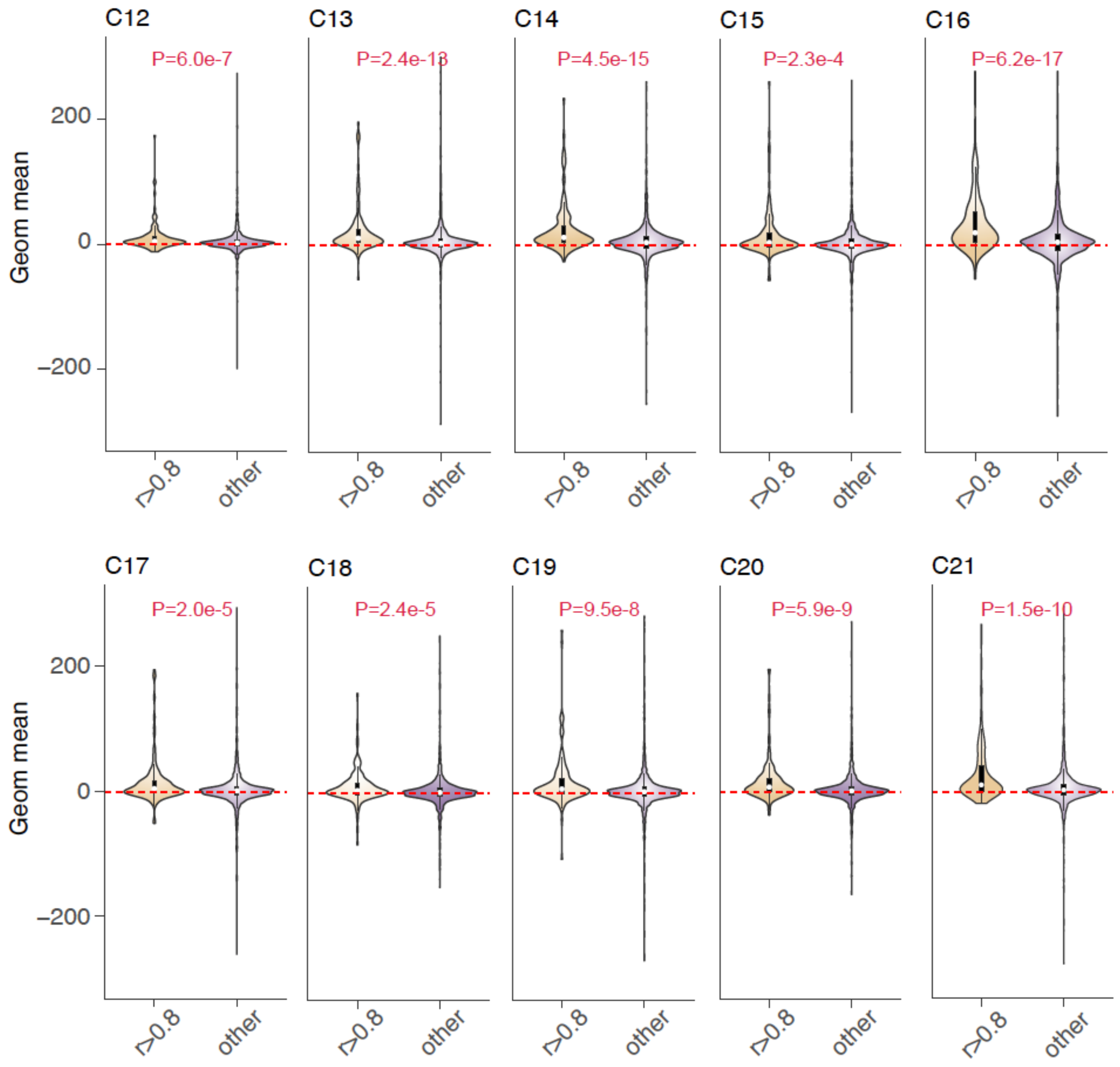


**Figure S4. Highly correlated mRNA-protein pairs tend to co-evolve at different nodes in the phylogenetical tree.**

The distribution of signed geometric means of mRNA- and protein-PIC values at 10 nodes for each gene with high mRNA-protein PIC correlations (Pearson’s R > 0.8) and other genes.

Supplementary Figure 5

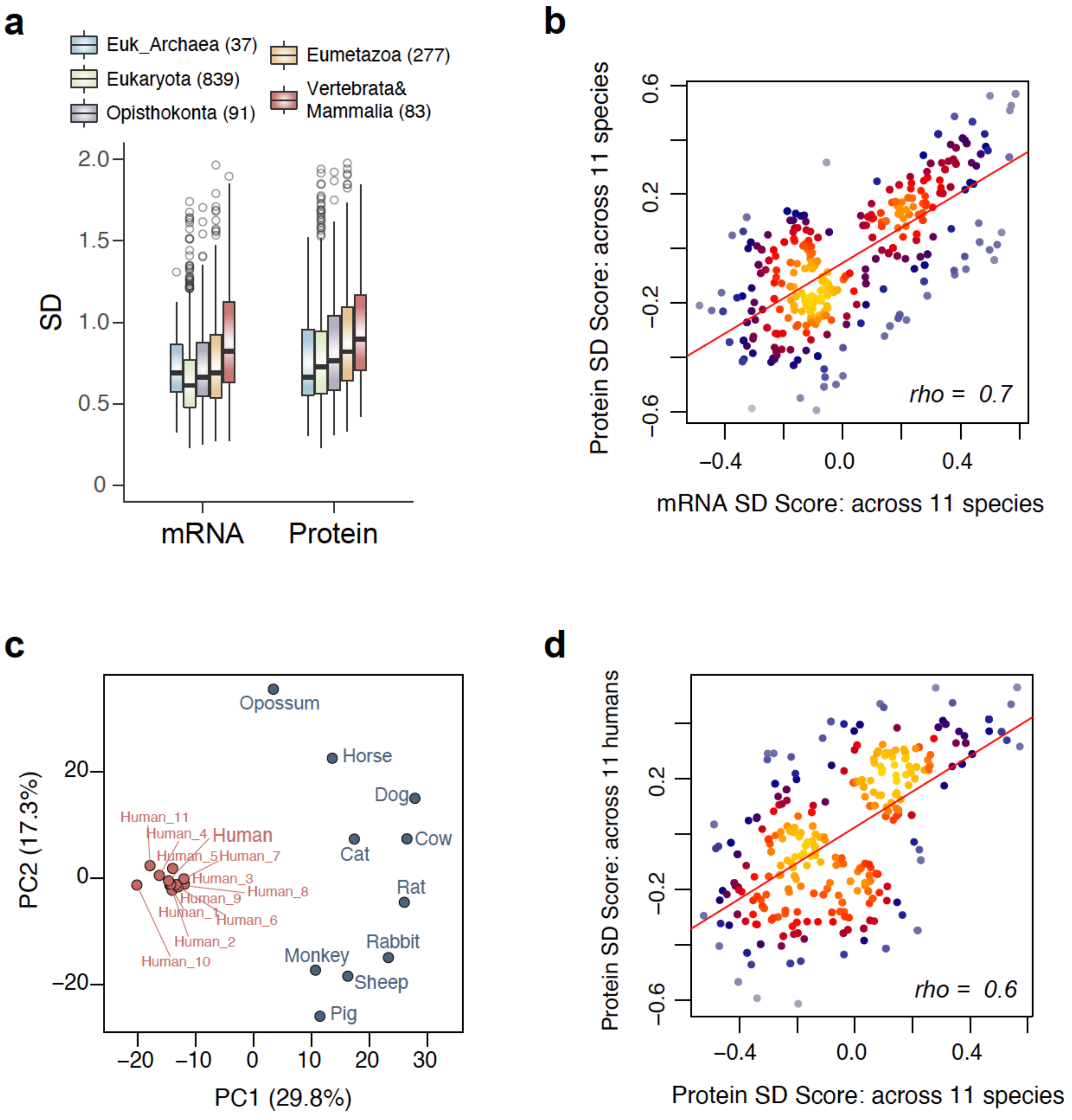


**Figure S5. Variability of mRNA and proteins and its functional relevance.**

(a) The newly evolved mammal-specific genes during evolution (i.e., "younger" genes) (Liebeskind et al., 2016) show higher expression divergence among mammals – especially at the protein level, than those "older" eukaryote-specific genes (b) All biological annotation items enriched by protein and mRNA SD across 11 species. Items of size >15 and P<0.05 are shown. (c) Principal component analysis of the proteome from 11 species and other 11 normal human individuals. The 11 individual human proteome data were also from skin fibroblast cells (Liu et al, 2017). (d) All biological annotation items enriched by inter-species and inter-individual protein SD. Items of size >15 and P<0.05 are shown.

Supplementary Figure 6

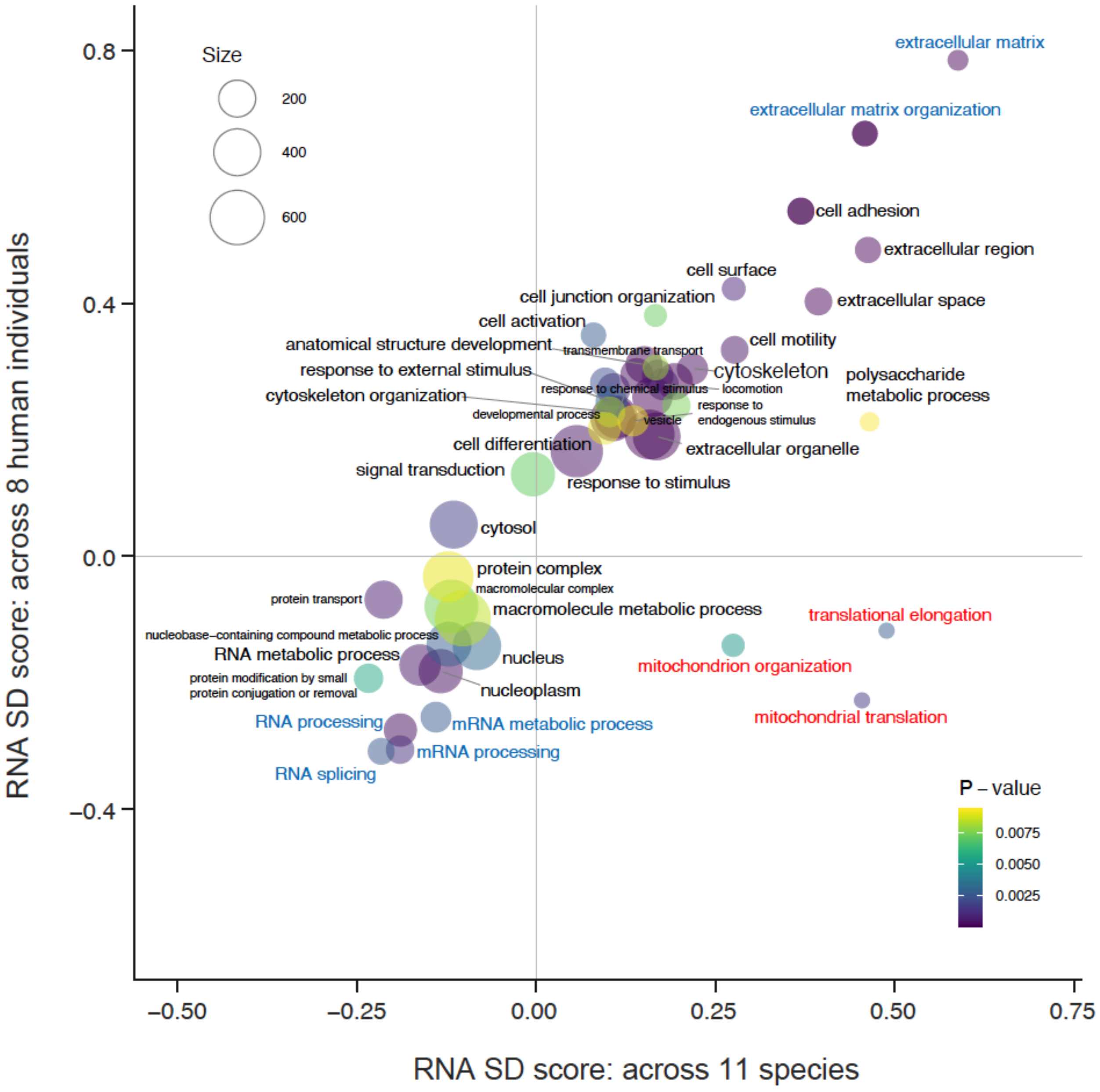


**Figure S6. RNA expression variation across species and human individuals.**

2D enrichment plot of significant GOBP slim or GOCC slim terms by SD (P <0.01). The axes denote enrichment score for the RNA SD from average across 11 species (x-axis) and 8 human individuals (y-axis).

Supplementary Figure 7

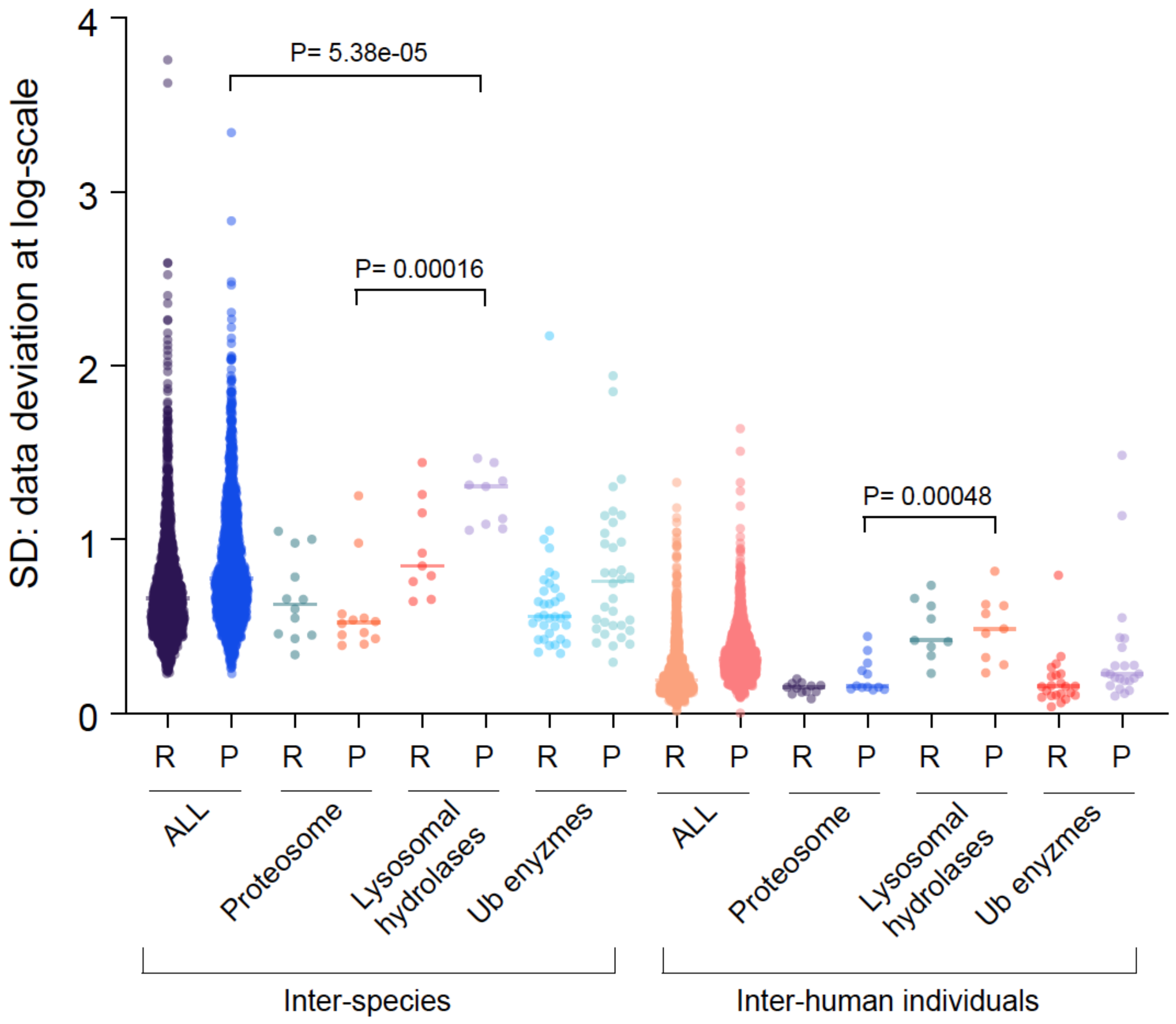


**Figure S7. The inter-individual and inter-species variability at the RNA and protein levels for protein degradation machinery.**

Note, the proteasome, lysosomal hydrolases and ubiquitin (Ub) enzymes including both deubiquitinating enzymes (DUBs) and E3 Ubiquitin ligases were summarized based on their mRNA and protein concentration variabilities, across species and across human individuals. R, mRNA; P, protein.

Supplementary Figure 8

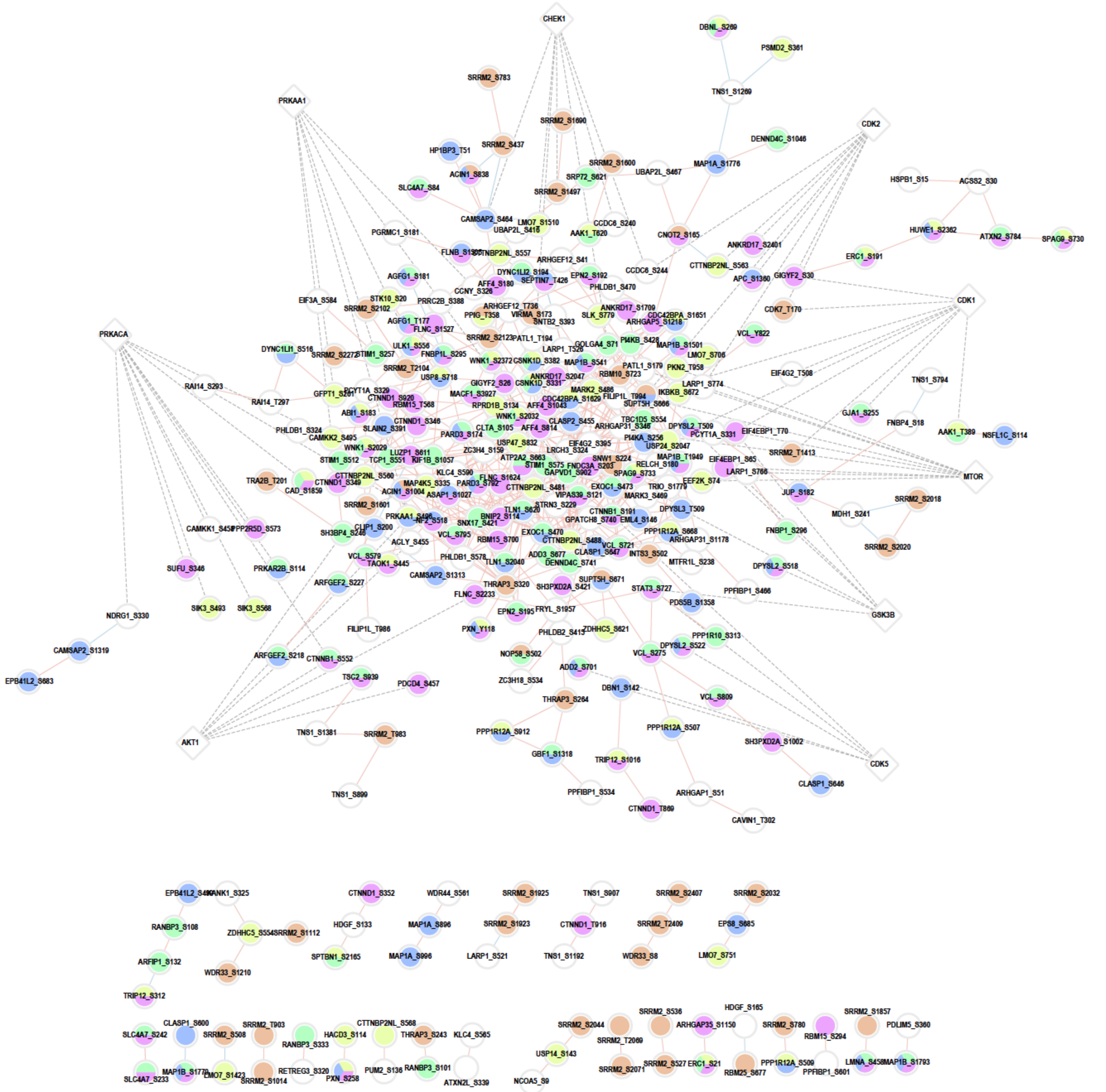


**Figure S8. Interaction network of phosphosites and the top kinases.**

The expanded version of the interaction network analysis of the significantly associated common P-sites (Pearson's correlation, $p < 0.001$) after correction by total protein changes. Red lines indicate positive associations while blue lines indicate negative associations. The representative GOBPs of phosphoproteins were highlighted in different colors. The top 9 kinases curated from the P-sites as substrates were shown as diamonds and the kinase-substrates pairs were shown as dashed lines.

Supplementary Figure 9

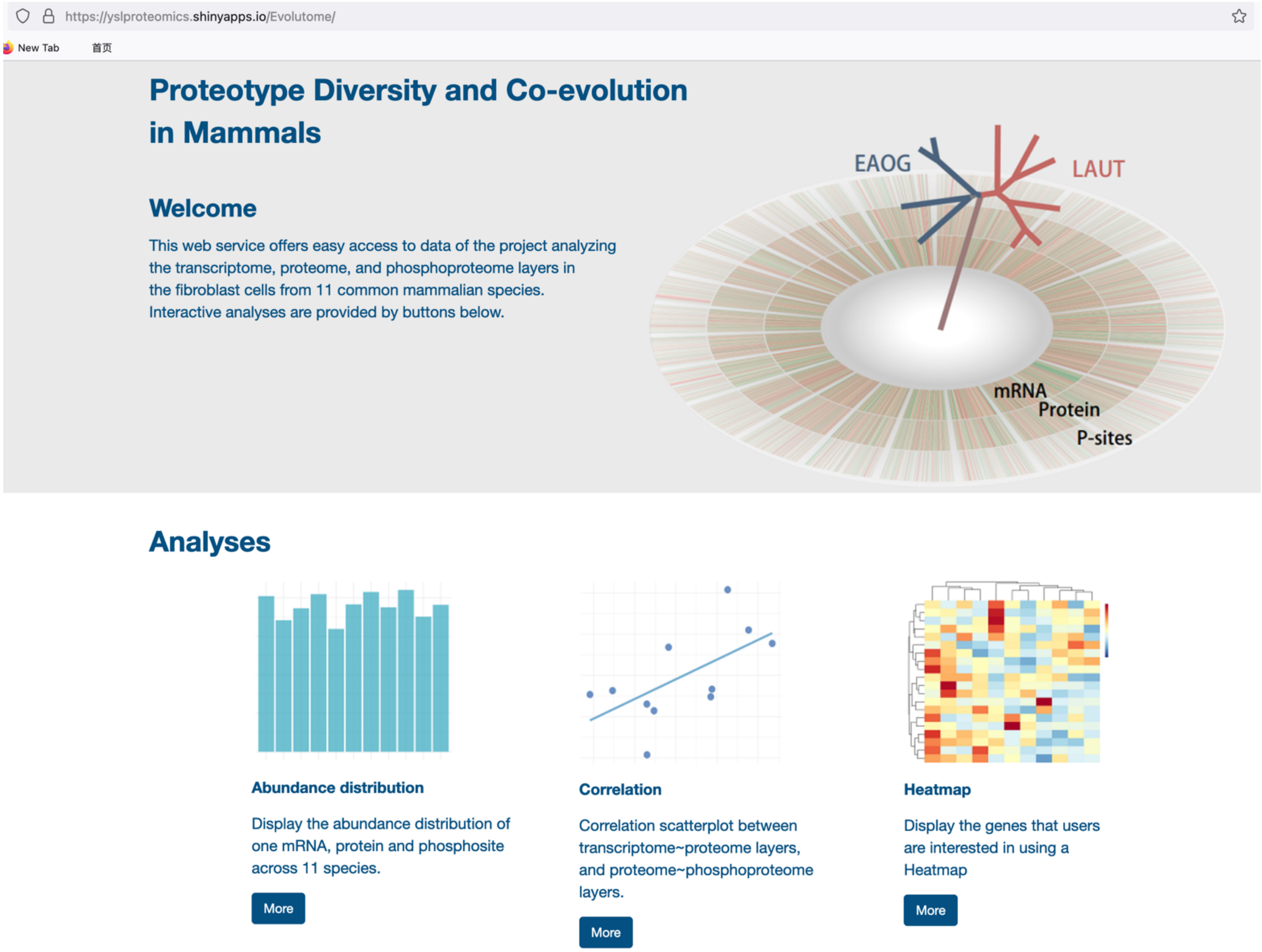


**Figure S9. A website to navigate and contemplate the data basis in 11 mammal species.**

This website interactively provides fast queries about the abundances for any transcript or protein in every species based on gene names. It additionally offers heatmap visualization and correlation scatterplots for genes or gene sets of interest between molecular layers.
